# Compensatory Mechanisms for Preserving Speech-in-Noise Comprehension Involve Prefrontal Cortex in Older Adults

**DOI:** 10.1101/2024.03.08.584193

**Authors:** Zhuoran Li, Yi Liu, Xinmiao Zhang, Nuonan Kou, Xiaoying Zhao, Xiangru Jiang, Andreas K. Engel, Dan Zhang, Shuo Wang

**Affiliations:** Department of Psychological and Cognitive Sciences, Tsinghua University, Beijing, China; Tsinghua Laboratory of Brain and Intelligence, Tsinghua University, Beijing, China; Department of Psychiatry, University of Iowa Carver College of Medicine, Iowa, United States; Stead Family Department of Pediatrics, University of Iowa Carver College of Medicine, Iowa, United States; Beijing Institute of Otolaryngology, Otolaryngology-Head and Neck Surgery, Beijing Tongren Hospital, Capital Medical University, Beijing, China; Department of Neurophysiology and Pathophysiology, University Medical Center Hamburg-Eppendorf, 20246 Hamburg, Germany

## Abstract

The capacity of comprehending others amidst noise is essential for human communication. However, it presents significant challenges for the elderly who often face progressive declines in the peripheral auditory system and the whole brain. While previous studies have suggested the existence of neural reserve and neural compensation as potential mechanisms for preserving cognitive abilities in aging, the specific mechanisms supporting speech-in-noise comprehension among the elderly remain unclear. To address this question, the present study employs an inter-brain neuroscience approach by analyzing the neural coupling between brain activities of older adults and those of speakers under noisy conditions. Results showed that the neural coupling encompassed more extensive brain regions of older listeners compared to young listeners, with a notable engagement of the prefrontal cortex. Moreover, the neural coupling from prefrontal cortex was coordinated with that from classical language-related regions. More importantly, as background noise increases, the older listener’s speech comprehension performance was more closely associated with the neural coupling from prefrontal cortex. Taken together, this study reveals the compensatory recruitment of neurocognitive resources, particularly within the prefrontal cortex, to facilitate speech processing in the aging brain, and further highlights the critical role of prefrontal cortex in maintaining the elderly’s ability to comprehend others in noisy environments. It supports the neural compensation hypothesis, extending the knowledge about the neural basis that underlies cognitive preservation in the aging population.

## Introduction

Understanding other people’s speech, commonly taking place in environments with background noise in daily life, poses a significant challenge for older adults. This challenge arises from the progressive decline in the auditory system’s structure and function (*1*), as well as alterations in the perisylvian language regions that are crucial for processing speech acoustics and mapping them into meaningful content (*2*). To counteract these age-related declines and maintain the cognitive ability in older adults, it is hypothesized that the aging brain may recruit additional neurocognitive resources (*3-5*). The neural reserve hypothesis suggests that the brain’s functional network possesses resilience and contains redundancy of neural resources (*6*), which allows it to maintain cognitive function despite age-related degradation or heightened demands imposed by difficult cognitive tasks (*3, 4*). Empirical evidence in support of this hypothesis includes a reserve and even upregulation of the neural activation of the language-related brain regions in older adults (*2, 7-9*), which may serve as a mechanism for preserving speech comprehension abilities. Alternatively, the neural compensation hypothesis proposes that when primary cognitive networks become less efficient, additional brain regions or circuits are recruited to sustain cognitive function (*3, 4, 10*). For instance, the prefrontal cortex, which shows minimal activation in younger adults (*2, 4, 11, 12*), has been reported to be significantly engaged during older adults’ speech processing. This compensation is hypothesized to be based on neural plasticity and functional reorganization of the brain, suggesting a dynamic adaptation to cognitive demands (*10*). However, despite these findings and hypotheses, the precise neural mechanisms that facilitate daily-life speech comprehension in older adults remain unclear.

A critical question concerns the extent to which various brain regions are engaged in understanding other’s speech in a naturalistic context. Despite significant efforts in exploring this question, most existing studies have focused on reduced speech conditions, using simple language materials like syllables or words. These experimental setups fail to replicate the complexities of real-world speech comprehension, which often involves more naturalistic speech, e.g., narratives, and is frequently impeded by environmental noise (*13, 14*). The processing of natural speech is more complex than that of simple stimuli, requiring more extensive computational analysis of meaningful content and context, thus engaging a wider network of fronto-parieto-temporal cortical regions (*13, 14*). Furthermore, adapting to background noise demands additional neurocognitive resources to mitigate the interference of the background noise and enhance selective processing of the target speech (*15-17*). Consequently, previous studies may have significantly underestimated the neurocognitive demands of processing natural speech in noisy conditions, resulting in an incomplete understanding of the brain regions involved in speech comprehension in the aging brain.

Another critical issue is the functional role of various brain regions in natural speech comprehension among older adults. While previous studies have noted enhanced activation in the prefrontal cortex and classical language regions (*2, 18*), the specific regions most influential to speech ability in older adults remain much controversial. The neural reserve hypothesis supposed that the preservation of older adult’s cognitive ability was achieved based on the functional reserve or reorganization within the original functional network (*3*). Following this line, the older adult’s speech performance should be closely related to the neural responses within the language network. In contrast, the neural compensation hypothesis emphasizes the recruitment of brain regions or networks that is not typically involved in young adults, e.g., prefrontal cortex, to facilitate the cognitive maintenance (*3, 19*). Drawing on this, the older adult’s speech ability may be more closely associated with the neural responses of non-language-specific regions, such as the prefrontal cortex, rather than the language-related regions. Considering the limited capacity of neural reserve and the overall decline of the language network within the aging brain (*3, 4*), we hypothesize that the mechanism of the neural compensation plays a more significant role in maintaining the speech performance or ability, especially under noisy conditions that demand more cognitive resources. Specifically, the prefrontal cortical response may be more predictive of the actual speech-in-noise performance in older adults. Notably, as the prefrontal cortex may facilitate speech processing by compensating for the language network rather than replacing it (*20, 21*), we further hypothesize an enhanced functional integration between them within the aging brain (*22, 23*) for facilitate the natural speech processing in older adults.

To address these questions and test our hypotheses, we conducted a study with a cohort of older adults aged 59-71 years, who listened to pre-recorded narratives spoken by young adults. To simulate real-life speech scenarios with environmental noise, we added white noise to the narrative audios at four different levels: no noise, +2 dB, -6 dB, and -9 dB. This design allowed us to explore how various brain regions adapt and function under different noise conditions. Neural activities from both the old adults and young speakers were recorded by functional near-infrared spectroscopy (fNIRS), chosen for its motion insensitivity and low operational noise than functional magnetic resonance imaging (*24*). To elucidate the involvement of different brain regions in natural speech processing, we adopted an inter-brain neuroscience approach (*25-27*). This approach analyzes the coupling between the neural activities of old adults and those of the speakers, with the latter providing a reference for understanding the neural response to speech. This method avoids the difficulty of disentangling the interdependent speech features (*13, 28*) and captures additional information beyond common speech features, such as acoustics and semantics (*29-33*). Based on this approach, we examined the speaker-listener neural coupling from various brain regions across different noise levels (*14, 34, 35*). Furthermore, we compared the neural coupling of old listeners to a control group of young listeners (*35*) to further highlight age-specific neural mechanisms.

Our findings reveal that the neural activity in the prefrontal cortex and language-related regions of old listeners was significantly coupled with the speaker’s brain activity, regardless of the noise level. These couplings involved a wider range of brain regions in old listeners compared to young listeners, suggesting both neural reserve and neural compensation mechanisms within the aging brain. Notably, with the increasing of background noise, the old listener’s comprehension performance was more closely associated with their neural coupling from the prefrontal cortex. This relationship demonstrates the functional role of the prefrontal cortex in compensating for reduced auditory input and to maintain speech comprehension, highlighting the functional role of neural compensation in preserving the speech-in-noise comprehension ability among the older adults. Moreover, the older listener’s neural coupling of prefrontal cortex was robustly correlated to that of the language regions across noise levels, which was not observed in the young listeners. These results suggest that the prefrontal cortex was closely integrated with the classical language network in the aging brain. Taken together, our study elucidates the neural mechanisms that support natural speech comprehension in noise among older adults. It highlights the critical role of the prefrontal cortex in the cognitive preservation of speech(-in-noise) comprehension ability, shedding lights on potential interventions to promote successful cognitive aging.

## Results

### Noise impacts old listener’s perception and speech comprehension

Figure 1A illustrates the experimental design of this study. A group of older adults with normal or close-to-normal hearing threshold were recruited to listen to narratives pre-recorded from a group of younger speakers (*35*). The listener’s experiment was conducted separately after the speaker’s experiment, for the convenience of controlling the speaking quality of speakers and incorporating noise into the speaker’s narrative audios (*33, 35, 36*). The neural activities from the speakers and the listeners were measured by fNIRS with the same coverage of brain regions, including prefrontal cortex and classical language regions (Figure 1B, see Methods for more details). The listeners’ subjective ratings of the narratives’ clarity and intelligibility and their comprehension scores were collected. Results are shown in Figure 2A. The repeated-measures analysis of variance (rmANOVA) revealed that noise level significantly affected the old listeners’ perception and comprehension to the speech (clarity: *F*(3, 84) = 76.17, *p* < .001, partial 77^2^ = .73; intelligibility: *F*(3, 84) = 51.15, *p* < .001, partial 77^2^ = .65; comprehension: *F*(3, 84) = 19.69, *p* < .001, partial 77^2^ = .41). Post hoc analysis showed that, for the clarity and the intelligibility ratings, all pairwise comparisons between noise levels were significant (post hoc *t*-test, *p*s < .001, corrected by false discovery rate (FDR), (*37*)), with a lower rating when noise increased. For the comprehension score, except for the comparison between NN and 2 dB (post hoc *t*-test, *p* > .999), the other pairwise comparisons were significant (post hoc *t*-test, *p*s < .05, FDR corrected). Notably, the comprehension scores of all the listeners are substantially higher that the random level (25%) in all conditions. Even in the strongest noise level (-9 dB), the average score was 65.3±2.6% (mean±SE), suggesting a pretty good preservation of speech-in-noise comprehension ability for the old listeners in this study.

**Fig. 1.**
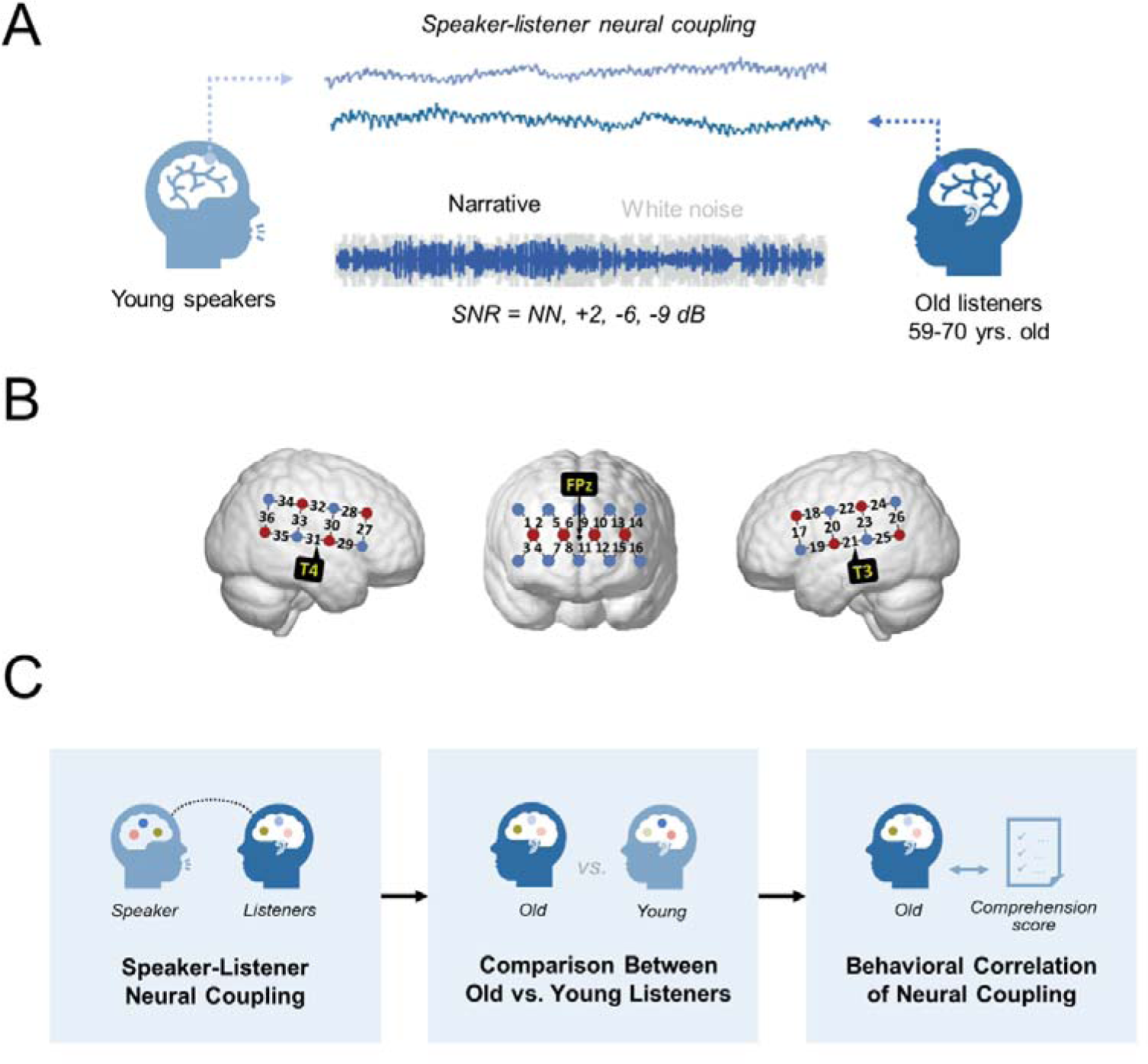
Experimental design and behavioral results. (A) Overview of the experimental design. (B) The fNIRS optode probe set. The red dots represent the source optode; the blue dots represent the detection optode. Channels of 21 and 31 were placed at T3 and T4, and the center of the prefrontal probe set was placed at FPz, in accordance with the international 10-20 system. The probe set covered the prefrontal cortex and classical language-related regions in both hemispheres. (C) Overview of the analysis framework. The inter-brain neural coupling between the speaker and old listener was analyzed first. The older listener’s neural coupling was then compared to that from a group of young listeners (*35*) to examine how different brain regions are compensatorily involved within the aging brain. Moreover, the functional role of these neural coupling was further exmained by analyzing its correlational relationship to the older listener’s speech comprehension performance in noisy conditions.

**Fig. 2.**
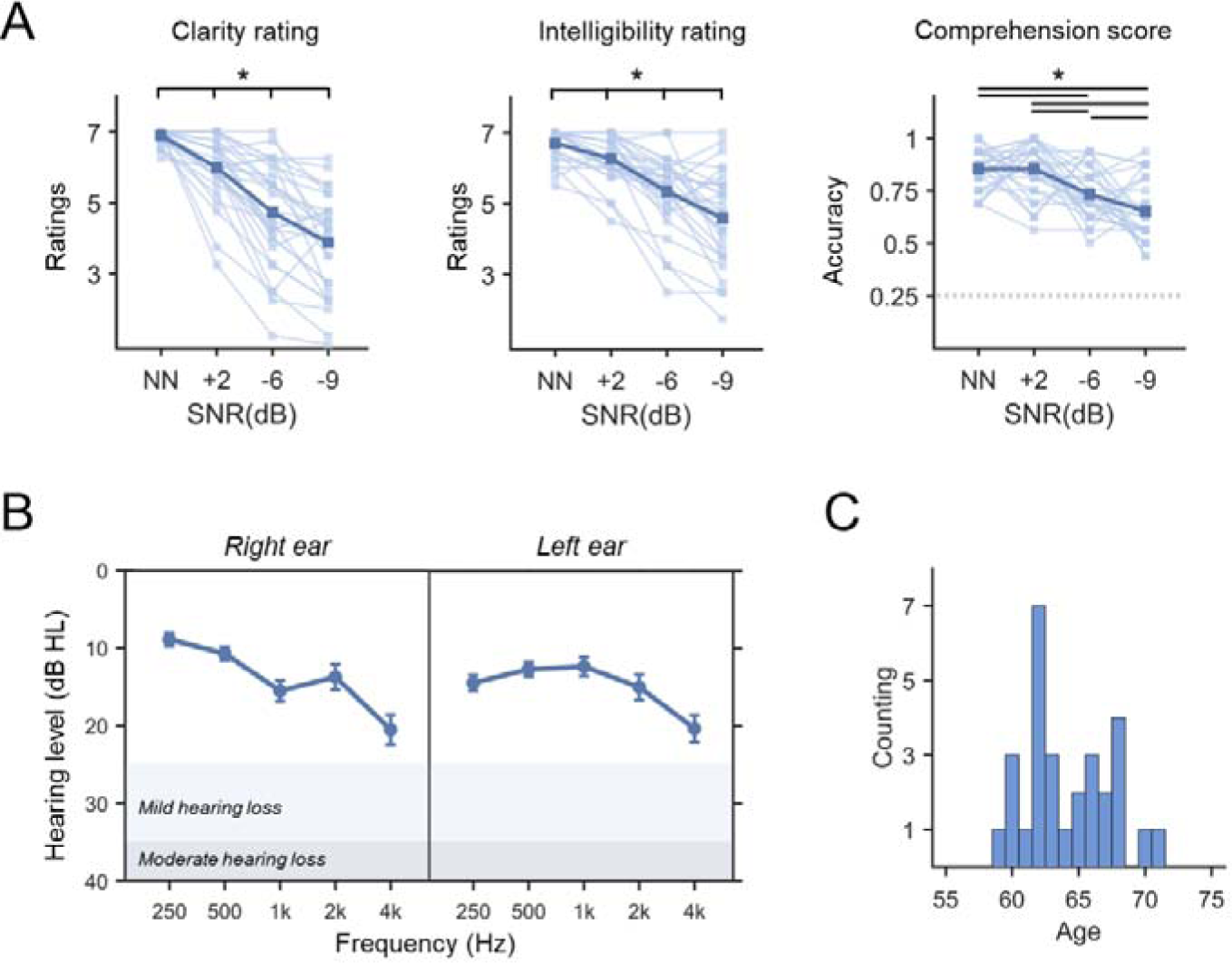
(A) Behavioral performance of old listeners. Results of repeated measures ANOVA showed that the noise significantly impacted the old listener’s clarity, intelligibility ratings and comprehension scores. Post hoc analysis revealed that for the clarity and intelligibility ratings all pairwise comparisons between four noise levels were significant. For the comprehension score, except for the comparison between NN and +2 dB, the other pairwise comparisons were significant. The asterisk (*) means significant result (*p* < .05, FDR corrected). (B) The average pure-tone hearing thresholds of the old listeners. The error bar means the standard error. (C) The age of the old listeners.

To examine the relationship between the perceptual ratings and comprehension performance, the correlation coefficients were computed across these behavioral measures. Results showed that the average ratings of clarity and intelligibility was significantly correlated (Spearman *r* = .40, *p* = .03). In contrast, the correlation between these two ratings and the comprehension score was not significant (*p*s > .23), suggesting a dissociation between the old listener’s subjective percept and the actual comprehension of speech. Besides, no significant correlation between these behavioral measures and the old listener’s hearing thresholds was found (*p*s > .14).

### Old listener’s neural activities are coupled to the speaker

To examine how old listener’s neural activities were coupled to the speaker, we conducted an rmANOVA on the speaker-listener neural coupling for all four noise levels and the resting-state baseline. This allowed to test for significant differences in neural coupling between any of the noise levels and the baseline as well as significant differences in coupling strength across different noise levels. A non-parametric cluster-based method (see Methods for more details) identified 57 clusters where speaker-listener neural coupling differed among the five conditions. These clusters covered the bilateral superior temporal gyrus (STG), middle temporal gyrus (MTG) (CH19/21/29/31) and right inferior frontal gyrus (IFG) (CH27) at the speaker’s side, which were likely to be speech-production-related brain regions (*32, 38*). At the listener’s side, the clusters covered an extensive range of brain regions, including the wide range of the prefrontal cortex (CH2, CH5-16) and the classical language regions. Based on the anatomical localization and functional organization within the language network (*35, 36, 39, 40*), these language regions were divided into ventral language regions, including right STG, MTG (CH29/31/35), bilateral inferior parietal lobe (IPL, CH23/24/26/33/34/36), and dorsal language regions, including bilateral IFG (CH17/27), bilateral pre- and post-central gyrus (pre-/postCG, CH18/20/22/28/30/31). The channel and localization of these channel combinations are shown in Figure 3A and S1A. The strength of the neural coupling for each cluster is shown in Figure S2. For all these clusters, the frequency band was around 0.01-0.03 Hz, as shown in Figure S1B. It was in the same frequency band as discovered in our previous studies on young listeners (*35, 36*). The specific information of all the clusters, including the channels of speaker and listener, frequency band, and cluster-level statistics, are listed in Table S1.

**Fig. 3.**
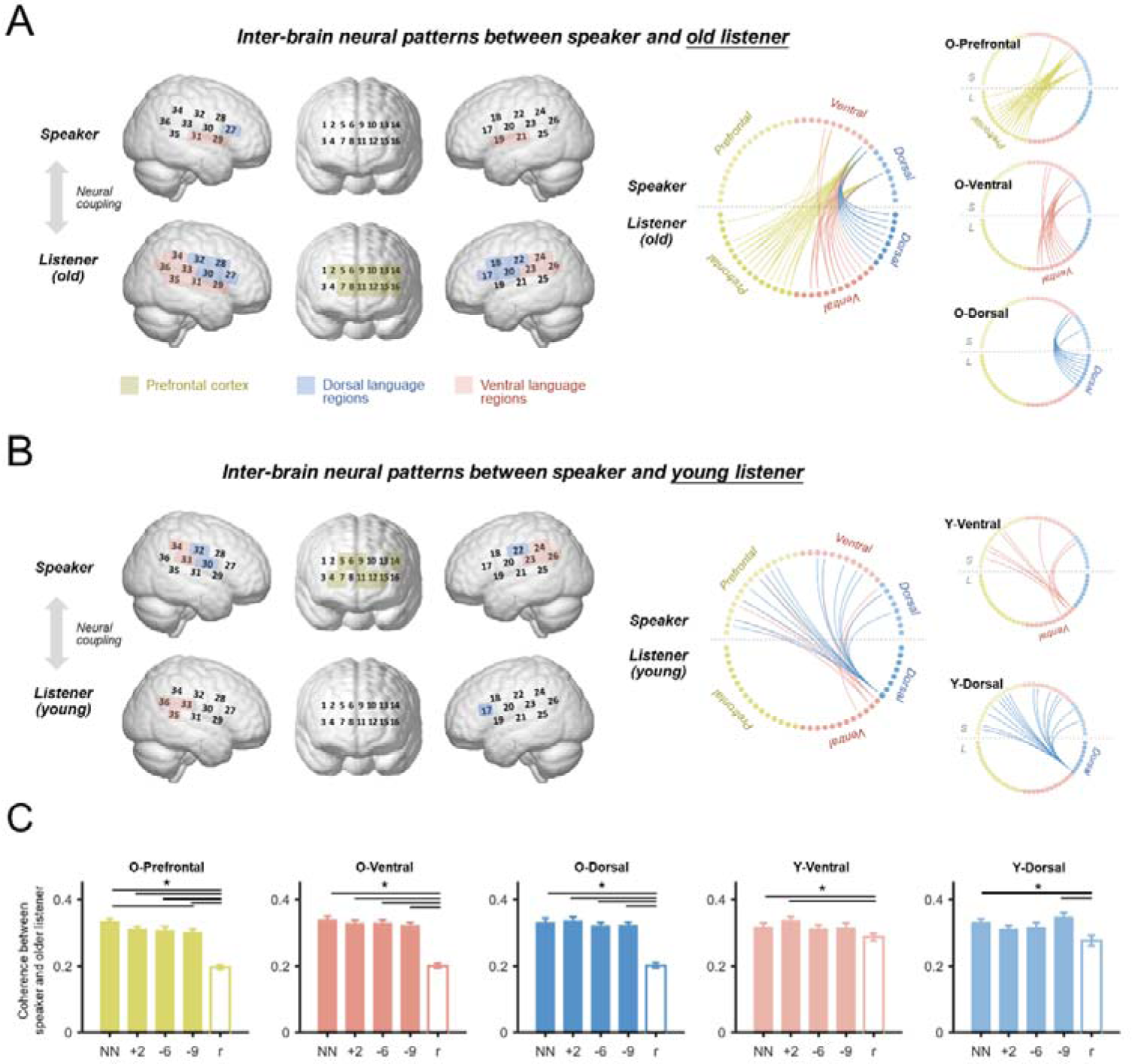
Speaker-listener neural coupling. (A) Significant speaker-listener neural coupling for the old listeners. The old listener’s neural activities from prefrontal cortex, bilateral ventral and dorsal language regions were coupled to the speaker’s neural activities. Different color means different anatomical brain regions: Yellow means the prefrontal cortex; red means the ventral language regions; blue means the dorsal language regions. Based on the brain region at the listener’s side, these inter-brain neural coupling were divided into three patterns: O-Prefrontal, O-Ventral, and O-Dorsal. (B) The inter-brain neural patterns for the young listeners. They were divided into two patterns: Y-Ventral, Y-Dorsal. (C) The old listener’s speaker-listener neural coupling from five inter-brain neural patterns. For O-Prefrontal, O-Ventral and O-Dorsal, the neural coupling in all the four noise levels was higher than the resting-state baseline. Besides, for the O-Prefrontal pattern, the coupling in NN was higher than -9 dB. For the Y-Ventral pattern, the coupling at NN and +2 dB was higher than that at the baseline; for the Y-Dorsal pattern, the coupling at NN and -9 dB was higher than that at the baseline. The error bar means the standard error. The lines and asterisks (*) indicate significant results (*p* < .05, FDR corrected).

Based on the brain region at the listener’s side, these inter-brain clusters were grouped into three patterns (Figure 3A), i.e., O-Prefrontal (prefrontal cortex, “O” refers to clusters from the old listeners), O-Ventral (ventral language regions) and O-Dorsal (dorsal language regions). The neural coupling for these three inter-brain patterns was derived by averaging the coupling values across clusters within the same group. These results at the pattern level are illustrated in Figure 3C. Post hoc analysis showed that for all of these three patterns, the neural coupling at the four noise levels were significantly higher than the resting-state condition (post hoc *t*-test, *p*s < .05, FDR corrected), indicating that the neural activities of old listeners were significantly coupled to the speaker in both quiet and noisy conditions. Besides, the neural coupling of the O-Prefrontal pattern at NN is higher than -9 dB (post hoc *t*-test, *p* = .016, FDR corrected), and marginally higher than that at 2 dB and -6 dB (post hoc *t*-test, *p*s = .052, .052, FDR corrected), suggesting its sensitivity to the appearance of background noise. However, no significant difference was observed between the neural coupling of the O-Prefrontal pattern and that from the O-Ventral or O-Dorsal patterns, even in the noisy conditions (*p*s > .10, FDR corrected).

### Distinct inter-brain neural patterns between old listeners and young listeners

To explore how age influenced the neural mechanisms of speech-in-noise processing, we compared the speaker-listener neural coupling of the old listeners to a group of young participants (Figure 3B) who listened to the same narratives (*35*). As illustrated in Figure 3A/B, the inter-brain neural patterns of the young listeners differed from that of the old listeners. As compared to the old listeners, the inter-brain neural patterns of the young listeners did not engage the prefrontal cortex, and only covered a few brain regions in the ventral language regions (CH33/35/36, right MTG/IPL) and the dorsal language regions (CH17, left IFG). In contrast, they covered extensive brain regions at the speaker’s side, involving the prefrontal cortex and a broad coverage of ventral and dorsal language regions. These clusters were also grouped into two patterns and named as Y-Ventral and Y-Dorsal, respectively (“Y” refers to clusters from young listeners, to distinguish them from the patterns of the old listeners), based on their anatomical location at the listener’s side. The information of the young listeners’ clusters is listed in Table S2.

We also examined whether the neural coupling from the Y-Ventral and Y-Dorsal patterns was significant in the old listeners (Figure 3C). Results showed that for the Y-Ventral pattern, the neural coupling at NN and +2 dB was higher than that of the resting-state (Paired samples *t*-test, *t*s = 2.25, 3.48; *p*s = .032, .005, FDR corrected; Cohen’s *d*s = 0.42, 0.65), -6 dB and -9 dB was marginally higher than the resting-state (Paired samples *t*-test, *t*s = 1.77, 1.57; *p*s = .054, .063, FDR corrected; Cohen’s *d*s = 0.33, 0.29). For the Y-Dorsal pattern, the neural coupling at NN and -9 dB was higher than that of the resting-state (Paired samples *t*-test, *t*s = 3.16, 3.17; *p*s = .005, .005, FDR corrected; Cohen’s *d*s = 0.59, 0.59), +2 dB and -6 dB were marginally higher than resting-state (Paired samples *t*-test, *t*s = 1.83, 1.73; *p*s = .054, .054, FDR corrected; Cohen’s *d*s = 0.34, 0.32). For the young listeners, the significance of neural coupling at O-Prefrontal, O-Ventral and O-Dorsal patterns was also examined. As shown in Figure S3, their neural coupling in all the four noise levels was significantly higher than that of the resting-state baseline (Paired samples *t*-test, *t*s > 3.00; *p*s < .05, FDR corrected; Cohen’s *d*s > 0.91).

To examine the specificity of these neural patterns for particular listener groups, we compared the strength of these inter-brain neural coupling between the old and the young listeners. The coherence of the resting-state baseline was first subtracted from that of the four noise levels, and then devoted to a two-way mixed-design ANOVA (between-participant factor: listener group/age; within-participant factor: SNR) to analyze how age impacted the neural coupling. Results showed that, for all the five patterns, the main effect of listener group was significant (*F*s > 4.11, *p*s < .05, FDR corrected). For the O-Prefrontal, O-Ventral and O-Dorsal patterns, the neural coupling of the old listener was significantly higher than that of young listener. In contrast, for the Y-Ventral and Y-Dorsal patterns, the neural coupling of the young listener was significantly higher than that of the old listener. The main effect of SNR and the interaction effect between the listener group and SNR were not significant (*F*s < 1.84, *p*s > .14). The group comparisons of the neural coupling are shown in Figure S4.

Furthermore, we compared the inter-brain neural coupling between two types of inter-brain neural patterns (O-Prefrontal/Ventral/Dorsal vs. Y-Ventral/Dorsal). A two-way rmANOVA (within-participant factors: inter-brain pattern, SNR) was conducted for each listener group. Results showed that the main effect of the inter-brain pattern was significant for both listener groups (old listener: *F*(4, 112) = 16.46, *p* < .001; young listener: *F*(4, 56) = 4.51, *p* = .003). Post hoc analysis further showed that, for the old listeners, the neural coupling from the O-Prefrontal, O-Ventral and O-Dorsal patterns was significantly higher than that from both the Y-Ventral and Y-Dorsal patterns (post hoc *t*-test, *p*s < .05, FDR corrected). In contrast, for the young listeners, the neural coupling from the O-Prefrontal, O-Ventral and O-Dorsal patterns was significantly lower than that from the Y-Dorsal pattern (post hoc *t*-test, *p*s = .040, .003, .007, FDR corrected), but not the Y-Ventral pattern (post hoc *t*-test, *p*s > .05, FDR corrected). The main effect of SNR and interaction effect between inter-brain pattern and SNR were not significant (*F*s < 1.56, uncorrected *p*s > .10). These findings suggested distinct inter-brain neural patterns for speech(-in-noise) processing in the old versus young listeners.

### Inter-brain neural coupling from prefrontal cortex correlates with old listener’s comprehension under noisy conditions

To examine the behavioral relevance of the speaker-listener neural coupling, a correlation analysis was performed between the old listeners’ speech comprehension scores and their neural coupling. The analysis was conducted both at the cluster level (Figure 4A) and at the pattern level (Figure 4B). At the cluster level, these *r*-values were computed for each cluster, and then Fisher-*z* transformed. These *r*-values were then devoted to a one-sample *t*-test to examine their significance in each noise level, and further compared between different noise levels to examine the effects of noise. At the pattern level, the correlation between the average neural coupling of all clusters for one particular inter-brain neural pattern and comprehension score was computed. For the O-Prefrontal pattern, significant correlation was observed in the moderate or strong noise level. At the cluster level, the *r*-values were significant at -6 dB and -9 dB (One sample *t*-test, *t*s = 5.84, 4.14; *p*s < .001, FDR corrected; Cohen’s *d*s = 1.12, 0.80), but not at NN and +2 dB (*p*s > .26). The pairwise *t*-test further showed that the *r*-values at NN were lower than those at the other three conditions (Paired samples *t*-test, *t*s < -3.21; *p*s < .05, FDR corrected; Cohen’s *d*s > 0.61), and the *r*-values at +2 dB were lower than those at -6 dB and -9 dB (Paired samples *t*-test, *t*s = -2.60, -2.06; *p*s = .023, .059, FDR corrected; Cohen’s *d*s = 0.50, 0.40). At the pattern level, the correlation between the neural coupling of the O-Prefrontal pattern and the comprehension score was significant at -6 dB and -9 dB (Spearman *r*s = .39, .37; *p*s = .035, .047, uncorrected), but not at NN and +2 dB (*p*s > .39).

**Fig. 4.**
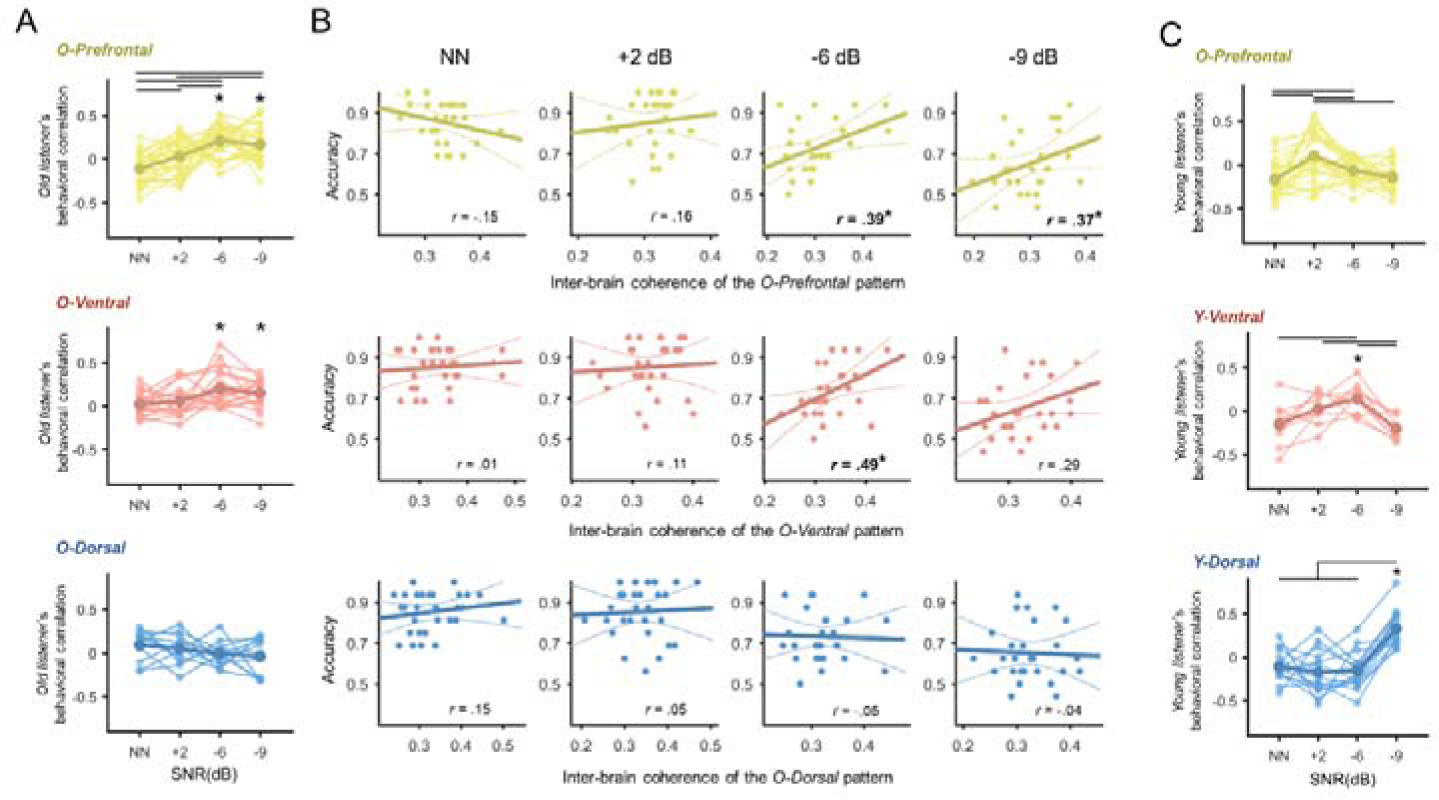
Behavioral relevance of speaker-listener neural coupling. (A) Correlation between the old listener’s comprehension score and the coupling from the clusters. Each line represents the correlation r-values (Fisher-z transformed) of one cluster belonging to one particular inter-brain neural pattern. The asterisk indicates significant r-values in one particular noise level; the line indicates a significant pairwise comparison between two noise levels (*p* < .05, FDR corrected). For both the O-Prefrontal and O-Ventral patterns, the *r*-values in -6 dB and -9 dB were significant. For the O-Prefrontal pattern, the *r*-values were higher when noise increased. (B) Correlation between the old listener’s comprehension score and average coupling for the inter-brain neural patterns. The neural coupling of the O-Prefrontal pattern was positively correlated with the comprehension score at -6 dB and -9 dB (Spearman *r*s = .39, .37; *p*s = .035, .047, uncorrected); the neural coupling of the O-Ventral pattern was correlated with the comprehension score at -6 dB (Spearman *r* = .49, *p* = .007, uncorrected). The other correlations were not significant. (C) Correlation between the young listener’s comprehension score and the couplings. For the O-Prefrontal pattern, the *r*-values under four noise levels were not significant, although significant differences existed across the different noise levels. For the Y-Ventral pattern, the *r*-values in -6 dB were significant (*p* < .05, FDR corrected); for the Y-Dorsal pattern, the *r*-values in -9 dB were significant (*p* < .05, FDR corrected).

For the O-Ventral pattern, significant correlation results were mainly observed in the moderate noise level. At the cluster level, the *r*-values were significant at -6 dB and -9 dB (One sample *t*-test, *t*s = 3.92, 3.89; *p*s = .001, .001, FDR corrected; Cohen’s *d*s = 0.92, 0.91), but not at NN and +2 dB (*p*s > .13). However, no significant results were found when comparing the results between any noise levels (*p*s > .07, FDR corrected). At the pattern level, the correlational relationship between the average neural coupling of the O-Ventral pattern and comprehension was only significant at -6 dB (Spearman *r* = .49, *p* = .007, uncorrected), but not in the other three conditions (*p*s > .12).

For the O-Dorsal and Y-Ventral patterns, no significant results were found at the cluster level (*p*s > .10, FDR corrected) or at the pattern level (*p*s > .10). For the Y-Dorsal pattern, at the cluster level, the *r*-values at NN and -6 dB were higher than those at +2 dB and -9 dB (Paired samples *t*-test, *t*s > 2.48; *p*s < .05, FDR corrected; Cohen’s *d*s > 0.64). However, these *r*-values were neither significant at the cluster level (*p*s > .07, FDR corrected) or at the pattern level (*p*s > .10).

To examine the functional role of the prefrontal cortex in younger listeners, the correlation between the younger listeners’ neural coupling from the O-Prefrontal pattern and speech comprehension scores were also analyzed (Figure 4C). At the cluster level, *r*-values at +2 dB were higher than those at the other three conditions (Paired samples *t*-test, *t*s > -2.55; *p*s < .05, FDR corrected; Cohen’s *d*s > 0.66), and *r*-values at NN were lower than -6 dB (Paired samples *t*-test, *t* = -3.21; *p* = .025, FDR corrected; Cohen’s *d* = 0.49). However, these correlations were neither significant at the cluster level (*p*s > .11, FDR corrected) or at the pattern level (*p*s > .29). In contrast, the young listener’s speech comprehension score was associated with the neural coupling from the Y-Dorsal at both the cluster level and the pattern level (*35*), as shown in Figure 4C.

### Neural coupling from prefrontal cortex integrates with language regions in old listener

To figure out how the prefrontal cortex was functionally connected with the language regions within the aging brain, the Spearman correlation between the neural coupling from these regions was computed under various noise levels and for both the old versus young listeners. For the old listeners, the inter-brain neural patterns of the O-Prefrontal, O-Ventral and O-Dorsal patterns were interconnected. At all the four noise levels, the neural coupling from the O-Prefrontal pattern was positively correlated with that from the O-Ventral pattern (Spearman *r*s = 45, .48, .38, .58; *p*s

= .014, .008, .043, .001, FDR corrected), and the O-Dorsal pattern (Spearman *r*s = 50, .51, .43, .38; *p*s = .006, .006, .022, .042, FDR corrected). The coupling strength from the O-Ventral pattern was also positively correlated with the that from the O-Dorsal pattern in all the four noise levels (Spearman *r*s = 76, .69, .46, .43; *p*s < .001, < .001, .013, .021, FDR corrected). These results suggested a close within-network connectivity of the language regions and the possible integration of the prefrontal cortex into the language network in older adults (Figure 5A/B).

**Fig. 5.**
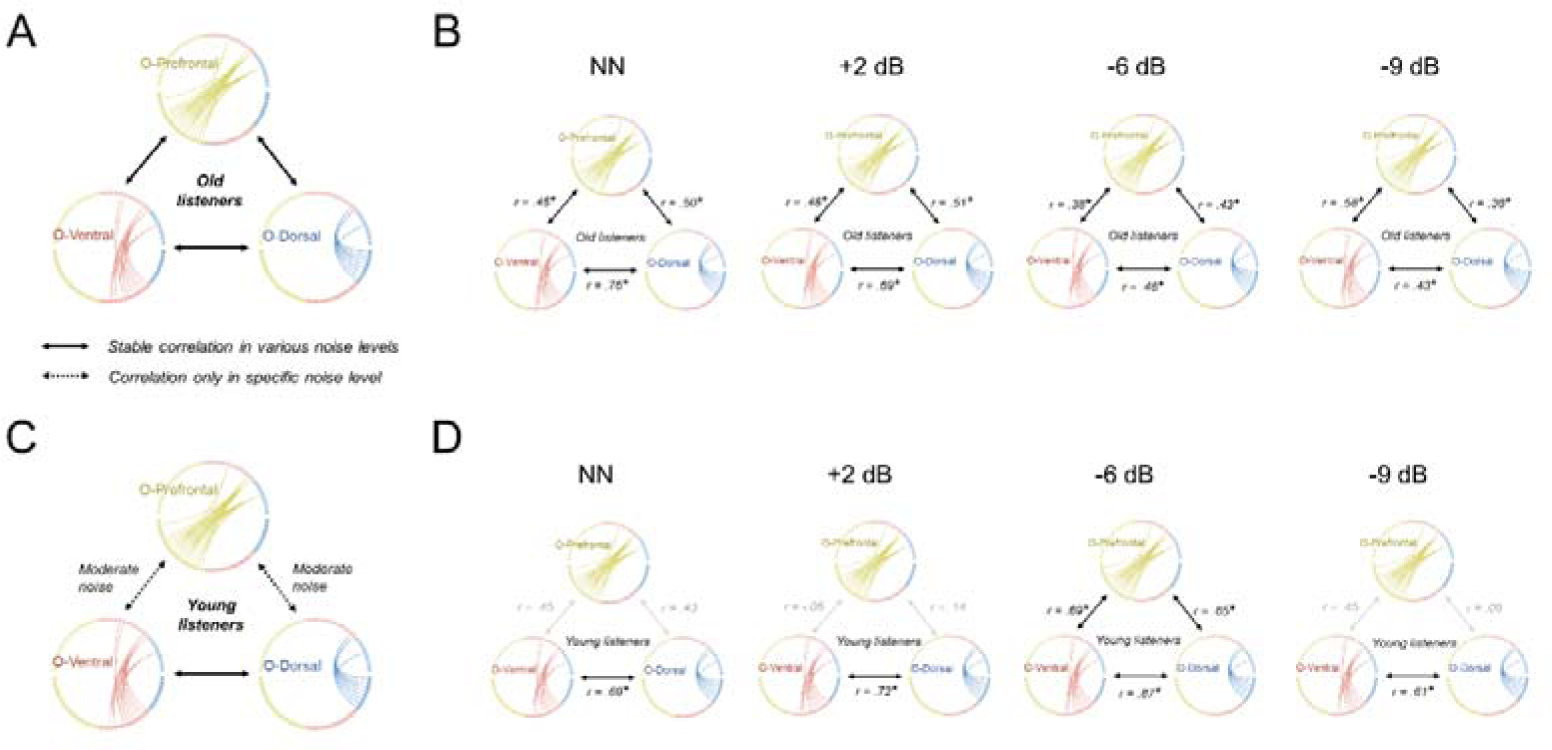
Integration between inter-brain neural patterns. (A/C) Summary of the integration between the inter-brain neural patterns of O-Prefrontal, O-Ventral and O-Dorsal for the old listeners (A) and the young listeners (C). (B/D) The correlation between three inter-brain neural patterns in four noise levels. Significant correlation were marked by the dark line and the asterisks (*p* < .05, FDR corrected). For old listeners, the integration between the O-Frontal pattern and the O-Ventral/O-Dorsal patterns was robust across noise levels. However, they were only observed in moderate noise level (-6 dB) for the young listeners. The O-Ventral and the O-Dorsal patterns was closely integrated in all four noise levels, for both the old listeners and the young listeners.

**Fig. 6.**
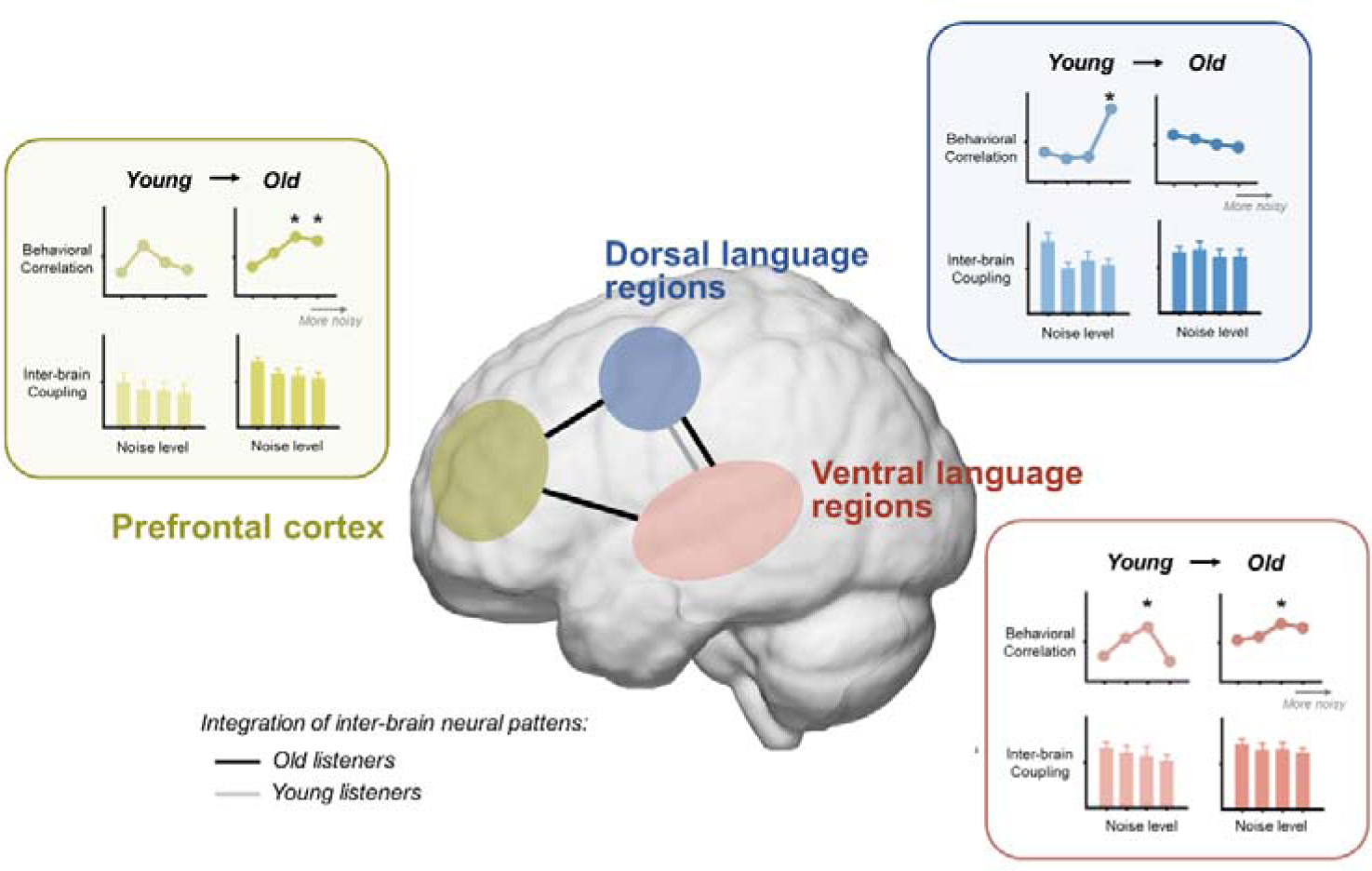
Summary of the neural mechanism for natural speech comprehension in different noise levels and age groups. In young adults, the dorsal and ventral language regions are engaged, with stable integration facilitating speech-in-noise processing. As noise intensity increases, the dorsal language regions play a more critical role, as indicated by a stronger correlation between the speech-in-noise comprehension performance and neural coupling. In contrast, older adults exhibit a compensatory recruitment of the prefrontal cortex to maintain speech comprehension abilities. This engagement is characterized by not only robust prefrontal cortex involvement across noise levels but also by an enhanced integration with language regions, a phenomenon not observed in young adults. Furthermore, the older adult’s speech-in-noise comprehension is more closely associated with the prefrontal cortex, especially as noise level increases. Asterisks (*) indicate significant correlational relationship between the speech comprehension performance and the neural coupling from the particular brain regions.

For the young listeners, the neural coupling from the O-Ventral and the O-Dorsal patterns was also correlated at all the four noise levels (Spearman *r*s = .69, .72, .87, .61; *p*s = .006, .004, < .001, .018, FDR corrected). However, the neural coupling from the O-Prefrontal pattern was only correlated with that from the O-Ventral and O-Dorsal patterns at -6 dB (Spearman *r*s = .65, .87; *p*s = .005, .010, FDR corrected), but not at the other noise levels (*p*s > .05, FDR corrected). These results indicated the prefrontal cortex might be less connected to the classical language regions for the young listeners. Interestingly, for both the old listeners or the young listeners, the correlation between the neural coupling from the Y-Ventral and the Y-Dorsal patterns were not significant at any noise levels (*p*s > .05, FDR corrected).

Besides, we also observed some correlation across two types of inter-brain neural patterns (Figure S5). For the old listeners, the neural coupling from the Y-Ventral pattern was positively correlated with that from the O-Prefrontal pattern at +2 dB (Spearman *r* = .44, *p* = .048, FDR corrected), and the O-Ventral pattern at NN and +2 dB (Spearman *r*s = .37, .44; *p*s = .017, .017, FDR corrected). The neural coupling from the O-Dorsal was negatively correlated with that from the Y-Dorsal pattern (Spearman *r* = -.51, *p* = .006, FDR corrected). For the young listeners, the neural coupling from the Y-Ventral pattern was correlated with that from the O-Ventral pattern at NN and -9 dB (Spearman *r*s = .53, .54; *p*s = .045, .042, FDR corrected), and the O-Dorsal pattern at -6 dB (Spearman *r* = .60, *p* = .021, FDR corrected). The neural coupling from the Y-Dorsal pattern was correlated with that from both the O-Prefrontal and O-Ventral patterns at NN (Spearman *r*s = .57, .54; *p*s = .030, .042, FDR corrected). These results might reflect a complex relationship between the two types of the inter-brain neural patterns.

### Single-brain activation from prefrontal cortex correlates with old listener’s speech-in-noise comprehension

To have a comprehensive understand the mechanism and function of the prefrontal cortex and language-related regions within the aging brain, a single-brain neural activation analysis was performed in addition to the inter-brain analyses. As shown in Figure S6, significant activation was observed in both quiet and noisy conditions. The prefrontal cortex was significantly activated at NN, +2 dB and -9 dB (*p*s < .001, < .001, = .001, FDR corrected), and marginally significant at -6 dB (*p* = .073, FDR corrected). The ventral and dorsal language regions were all significantly activated in all the four noise levels (*p*s < .05, FDR corrected). No difference was found when comparing the neural activations in difference noise levels (*p*s > .18). The neural activation of the old listeners was further compared to that of the young listeners (Figure S7). No significant result was observed in any brain regions and in any noise level (*p*s > .18, FDR corrected).

A correlation analysis was conducted to explore the relationship between the old listener’s brain activation and speech comprehension performance. As shown in Figure 5B, the activation of prefrontal cortex and ventral language regions were positively correlated with comprehension score at -9 dB (Spearman *r*s = .38, .38; *p*s = .041, .045, uncorrected). The correlations in the other noise levels and in the dorsal language regions were not significant (*p*s > .15). Given that both the inter-brain neural coupling and single-brain activation of prefrontal cortex correlated with the comprehension performance when noise became strong (-9 dB), we analyzed their correlational relationship to further examine whether a high coupling of the old listeners related to a high activation within the corresponding brain region. This correlation result was not significant (Spearman *r* = .03, *p* = .86). The correlation in various noise levels and from other brain regions are further shown in Figure S8.

## Discussion

This study utilized an inter-brain neuroscience approach to investigate the neural mechanisms of natural speech-in-noise comprehension among the older adults. Through an analysis of speaker-listener neural coupling, we discovered that the neural activities of old listeners from many brain regions in the prefrontal cortex and classical language networks were coupled with the neural activities of the speaker. Compared to the young listeners, the neural coupling of the old listeners engaged more extensive brain regions, particularly in the prefrontal cortex. Besides, the neural coupling from the prefrontal cortex was reliably correlated to that from the language-related regions across different noise levels, which was not observed in young listeners. These results imply that older adults engage and integrate more neurocognitive resources from the prefrontal cortex to facilitate their natural speech processing in noisy environments. More importantly, the old listener’s speech comprehension performance was more closely associated with the neural coupling in prefrontal cortex as background noise increased, highlighting the functional role of the prefrontal cortex for older adults to maintain their speech comprehension under more acoustically challenging scenarios. In conclusion, this study examined how the aging brain processed natural speech in noise, and highlighted the compensatory mechanism centered around the prefrontal cortex for supporting the cognitive preservation of speech ability in older adults.

This study elucidates how different brain regions are involved during speech-in-noise processing in older adults. Our findings showed that the old listeners’ neural activities across a wide range of brain regions and in both hemispheres were coupled to the speaker. Specifically, the older listeners not only recruited brain regions within the classical language networks but also additionally engaged the prefrontal cortex to facilitate speech processing. Furthermore, the neural coupling in the older listeners extended beyond the language-dominant hemisphere (i.e., the left hemisphere) to include homologous regions in the right hemisphere (*41-43*). It contrasts with the more localized neural coupling observed in young listeners, which only involved a few critical brain hubs in the language networks (*35, 36*). This evidence supports both the neural reserve hypothesis, which presumes the existence of resilience and resource redundancy within the primary functional network to adapt to aging or pathology (*3, 4, 6*), and neural compensation hypothesis, which proposes the involvement of additional brain regions or circuits for facilitating cognitive maintenance (*3, 4, 10*). Notably, the neural coupling observed in older listeners remained robust in both quiet and noisy conditions, indicating that the prefrontal cortex and language networks can consistently contribute to speech processing across diverse listening environments. Likewise, the older listeners in this study also exhibited a relatively stable comprehension performance in noisy conditions. The stability of both the neural coupling and the comprehension performance suggests that the neural mechanisms supporting speech processing in the aging brain may be resilient to varying levels of environmental noise.

More importantly, this study highlights the functional role of the prefrontal cortex in older adults. Although prior studies have widely reported the activation of the prefrontal cortex during speech processing in older adults (*2, 11, 22, 41*), its specific contribution to the preservation of speech functions remains controversial. Drawing on the neural compensation hypothesis, the older adult’s speech performance would be more closely associated with the neural response within the prefrontal cortex, particularly under noisy conditions with a heightened cognitive demand (*3, 4, 10*). Our findings provide the first empirical evidence for this hypothesis by showing that the speech comprehension performance of the old listeners was more closely correlated with the neural coupling from prefrontal cortex as the noise increased. Meanwhile, we also observed a significant correlation between the older adults’ speech performance and the neural coupling from the ventral language regions. However, it was only observed at moderate noise level, not at high noise level. This indicates that while the language network’s functionality is preserved in the aging brain, its adaptability is limited (*3, 4*). Consequently, older adults may rely on compensatory recruitment of resources beyond the language network to meet the cognitive challenges of increasing age or more complex tasks, such as increasing background noise intensity. Taken together, these findings highlight the importance of the neural compensation based on the prefrontal cortex for older adults to adapt to various noise conditions. Our results align with and extend existing theories that emphasize the compensatory role of the frontal lobe in successful aging, such as the Scaffolding Theory of Aging and Cognition (STAC) (*44, 45*) and the Posterior-Anterior Shift in Aging (PASA) (*46*), by demonstrating the compensatory role of the prefrontal cortex in natural speech(-in-noise) processing.

Our study also provides valuable insights into how the prefrontal cortex was coordinated for compensation with the classical language regions within the aging brain. By analyzing the correlation of speaker-listener neural couplings across various brain regions, we found that within the language network, coupling between ventral and dorsal language regions was consistently correlated across noise levels in both young and older adults. Notably, a robust correlation between the prefrontal cortex and the language network was observed exclusively in older adults, but not in young adults, suggesting an enhanced connectivity to facilitate speech processing in the aging brain. Our findings are consistent with existing discoveries of increasing global connectivity in older adults (*22, 47, 48*), which is thought to reflect a neural reorganization serving to offset the widespread neural deterioration (*48-50*). However, there explanation for this age-related increase in connectivity between the prefrontal cortex and the classical language network is controversial. The “interaction” view posits that the prefrontal cortex is involved in domain-general processing, such as inhibitory control and working memory, rather than language-specific functions, supported by evidence that its neural activity does not encode specific language or speech information (*20, 21*). Conversely, the “integration” perspective suggests that the prefrontal cortex acts as an extension of the language network following neural reorganization in the aging brain. Our study provides partial support for the integration perspective by showing that neural activities from both the prefrontal cortex and the language network are coupled to the same neural activities at the speaker’s side. This indicates that the prefrontal cortex may be involved in similar speech processing activities as the language network. Nevertheless, this debate remains unresolved and requires further research.

Interestingly, while some existing studies have found a relationship between the old listeners’ speech-in-noise ability and neural responses from the dorsal language regions, this relationship was not observed in our study. One possible explanation for this discrepancy may lie in the more naturalistic speech material used in our study. Most previous studies have used simplified language materials, such as syllables or words (*2, 11, 22, 41, 51*), which cannot fully capture how the brain processed natural speech, such as narratives or stories, in more realistic settings. In particular, the dorsal language regions could facilitate the speech-in-noise processing by generating an internal representation of speech (*52, 53*), e.g., enabling the clear phonological structures of the noise-masked speech by referring to pre-acquired phonological knowledge (*54*). Despite the essential role of dorsal language regions in identifying phonological units under noisy conditions (*9, 55*), the comprehension of natural speech is more closely associated with the processing of content and contextual information, especially for the older adults with a poor hearing ability (*56*). In this way, as found in our study, the natural speech-in-noise comprehension ability of older adults may be more closely related to brain regions responsible for processing meaning or content, which are predominantly located in ventral language regions as well as the prefrontal cortex, rather than the dorsal language regions (*57*). This discrepancy between our study and previous studies also highlights the importance of employing more naturalistic paradigms to understand the behavior, cognition and brain function of older adults in everyday settings.

It should be noted that neither the neural reserve nor neural compensation can fully restore the cognitive ability of older adults to the levels of young adults (*3, 4*). Actually, the additional cognitive or neural resources recruited reflect a reduced computational efficiency in the aging brain (*19, 58*). This study provides evidence for this from the inter-brain neural patterns observed on the speaker’s side. While the old listeners recruit a wide range of brain regions than the young listeners, their neural coupling at the speaker’s brain is limited to a smaller set of regions. Conversely, young listeners demonstrate neural coupling with a more extensive network of regions within the speaker’s brain, including bilateral temporo-parietal cortex and prefrontal cortex. The neural coupling between speakers and listeners from specific brain regions reflects the specific speech processing (*29, 59*). For example, coupling from the primary auditory cortex is associated with early acoustic processing (*32*), whereas coupling from higher-level regions like the posterior temporal lobe and frontal cortex is associated with more complex processes such as semantic and contextual interpretation (*29*). Therefore, given the inter-brain neural patterns showing limited coverage on the speaker’s side but extensive engagement of various brain regions on the listener’s side, our study indicates that older adults may have a diminished capacity for high-level speech processing and potentially low computational efficiency in the brain.

In addition to examining speaker-listener neural coupling, this study also analyzed neural activation in various brain regions. Both the prefrontal cortex and language-related regions exhibited significant activation across different noise levels. Importantly, similar to the findings from the neural coupling analysis, the neural activities from the prefrontal cortex and ventral language regions were positively correlated with speech comprehension performance. However, the correlation between the strength of speaker-listener neural coupling and neural activation was not significant, suggesting a dissociation between the specific speech processing within individual brain regions and the extent of their involvement. Neural coupling and neural activation capture speech-related neural responses in two different dimensions. Neural coupling focuses on the temporal dynamics of the listener’s neural activity, using the speaker’s brain as a reference, which may better capture the dynamic processing of natural speech. In contrast, neural activation emphasizes the amplitude of neural activity, which may reflect the engagement level of individual brain regions. Therefore, the analysis of speaker-listener neural coupling and single-brain neural activation can serve as complementary approaches (*33*). Future research could further explore the relationship between these two metrics and consider combining them to more fully elucidate the neural basis of speech processing in the brain.

In conclusion, the present study has revealed how natural speech is processed within the aging brain and has highlighted the compensatory role of the prefrontal cortex in preserving the speech-in-noise comprehension ability in older adults. Future studies could build upon our findings in the following aspects. Firstly, this study focused exclusively on healthy aging adults with normal hearing thresholds and good cognitive abilities. However, the old adult population exhibits significant heterogeneity across a broader age range (*60, 61*), regarding diverse hearing loss or impairments (*62, 63*), varying language experience (*64*), education backgrounds (*65*) and other traits. These variables can profoundly influence the structure, function, and (re)organization of the aging brain. To obtain a more comprehensive understanding of how the aging brain processes speech in noise, future studies could recruit different groups of older adults and further compare their neural responses during speech-in-noise comprehension. Secondly, this study identified and organized the speaker-listener neural coupling patterns based on the anatomical structure at the listener’s side. The recent development of advanced mathematical methodologies, e.g., dynamic inter-brain synchrony state analysis (*66*), offered a data-driven approach to define the inter-brain neural patterns. Future studies could adopt these techniques for a hypothesis-free exploration of inter-brain neural patterns (*67*). This could also help to better understand the functional organization of these brain regions. Thirdly, future studies could model speaker-listener neural coupling with multi-level features of speaker’s speech, e.g., acoustics and semantics, to further identify the computational processes underlying these compensatory inter-brain neural mechanisms. These speech features could be either modelled to the speaker’s or listener’s neural responses (*38*) or directly modelled to the temporal dynamics of speaker-listener neural coupling (*68*). This approach would provide further insights into the specific computational processing behind those age-related compensatory neural coupling. Lastly, while the present study indicated the compensatory role of the prefrontal cortex in older adults’ speech-in-noise comprehension ability, it remains to be explored whether this mechanism also supports or, conversely, hinders cognitive abilities in other domains. In particular, it has recently been proposed that neural resources may be deployed for enhancing the cognitive function in one area, e.g., the age-enhanced involvement of prefrontal cortex in speech-in-noise processing, reducing their availability for other cognitive functions *(69)*. Hence, to thoroughly examine the role of the prefrontal cortex in cognitive preservation (or decline) during age, future research might incorporate and analyze a broader spectrum of cognitive tasks. This would enable a more nuanced view of the cognitive and neural alterations in older adults.

## Methods

### Participants

Thirty older adults (15 males, 15 females; age 59 to 71 years old) participated in the study. The sample size was determined by a prior power analysis using G*Power (*70*), with the repeated measures *F*-test was employed (number of measurements = 4). The effect size was designed at the medium level of 0.25. The significance level (alpha) was set as 0.05, and the power was set as 0.9. All participants were native Chinese speakers, right-handed, and had an educational level equivalent to or higher than high school, ensuring basic reading proficiency. All participant passed the Chinese version of the Mini-Mental State Examination (MMSE) (> 26 scores). Based on the definition of hearing loss (World Report on Hearing by WHO, https://www.who.int/publications/i/item/9789240020481), one female participant was excluded due to a mild to moderate hearing loss as assessed by pure-tone audiometry (34 dB hearing level, average pure tone threshold between 500 and 4000 Hz). For the remaining twenty-nine participants, twenty-seven of them have a normal hearing threshold (≤20 dB), the rest two of them have a close-to-normal hearing threshold (22, 23 dB). Figure 2B shows the group average and standard deviation of their hearing levels.

The study was conducted in accordance with the Declaration of Helsinki and was approved by the local Ethics Committee of Tsinghua University and the Institutional Review Board of the Beijing Institute of Otolaryngology and Beijing Tongren Hospital. All participants provided written informed consent prior to the experiment.

### Experimental Design

The experimental design is illustrated in Figure 1A. The old listeners were instructed to listen to pre-recorded narrative audios under various noise conditions. These narratives were spoken by a group of young speakers, which were from our previous study (*35*). Both the neural activities of the old listeners and the young speakers were measured by fNIRS.

#### Stimuli

Each participant listened to sixteen narratives, with each lasting for 85 to 90 seconds. These narratives were about daily topics that were adapted from the National Mandarin Proficiency Test, such as favorite movies or travelling experiences. The narratives were spoken by four young adults (2 males, 2 females; 21 to 25 years old), all of whom were native Chinese speakers with professional training in broadcasting and hosting. Two of them (one male, one female) contributed five stories each, and two speakers (one male, one female) contributed three stories each. All narratives were about the speaker’s personal experiences, which were unfamiliar to the listeners. These audios were recorded by a regular microphone at a sampling rate of 44,100 Hz in a sound-attenuated room, and were then resampled to the 48,000 Hz to align with the speaker’s output frequency for this study.

Narrative audios were added with meaningless broadband white noise. For each listener, they were processed into one of four noise levels: no noise (NN) level and three noisy levels with the signal-to-noise ratio (SNR) of +2 dB, -6 dB, and -9 dB. Each noise level included four narratives. All processed audios were normalized for the average root mean square sound pressure level (SPL). For each narrative, four four-choice questions regarding narrative content were prepared to evaluate the listener’s comprehension performance. For example, one question was, “What is the relationship between the speaker and her friend? (••人和朋友是什••系)” and the four choices were 1) teacher-student, 2) cousins, 3) neighbors, and 4) roommates (1.•生•系, 2.姐妹•系, 3.•居•系, and 4.室友•系).

#### Experimental Procedure

The listener’s experiment was carried out in a sound-attenuated room. Sixteen narratives are structured into sixteen trials, with one narrative in each trial. During each trial, participants listened to one narrative under one of the four noise levels. After listening, they rated the clarity and intelligibility of the audio using a 7-point Likert scale, and completed a comprehension test based on the four choice questions. The comprehension score was calculated as the accuracy of these questions. Then, they could arrange a rest for a minimum of 10 seconds before proceeding to the next trial. One practice trial was conducted using an additional speech audio stimulus at -2 dB SNR level.

Before the task session, there is a 3-min resting-state session. Participants were instructed to get relaxed with their eyes closed. This resting-state session serves as the baseline for task session.

#### Neural measurement

The listeners’ neural activities were recorded from 36 channels using an fNIRS system (NirScan Inc., HuiChuang, Beijing). The fNIRS signals were collected at a sampling rate of 11 Hz, utilizing near-infrared light of two different wavelengths (730 and 850 nm). The concentration change of oxy-hemoglobin (HbO) and deoxy-hemoglobin (HbR) for each channel was then obtained based on the modified Beer– Lambert law.

Three sets of optode probes were positioned to cover the prefrontal cortex and bilateral inferior frontal, pre- and post-central, inferior parietal, middle and superior temporal gyrus, etc. As shown in Figure 1B, the positions of CH21 and CH31 were aligned with T3 and T4, respectively, as per the international 10-20 system. The central point of the prefrontal probe set was placed at the FPz position. To obtain a probabilistic reference to specific cortical region for each fNIRS channel, the topographic data was mapped onto a standardized MNI-space (Montreal Neurological Institute) (*71, 72*) by using NIRS_SPM (*73*). The MNI coordinates and anatomical labels for each channel are listed in Table S3.

### Neural data of speaker and young listeners

The neural data of the speakers and a group of young listeners were from our previous study (*35*). The speakers’ neural activities were recorded from the same coverage of brain regions, but by another fNIRS system (NirScan Inc., HuiChuang, Beijing) with a sampling rate of 12 Hz and using near-infrared light at three different wavelengths (785, 808, and 850 nm). These data were downsampled to 11 Hz to match the sampling rate of the old listener in this study.

The young listeners comprised fifteen young adults (7 males, 8 females; ages 18 to 24 years old). Their neural signals were obtained using the same fNIRS system, with the same brain coverage as the speakers. The experimental design for the young listeners were the same as that for old listeners in this study. However, they listened to a larger number of narratives (thirty-two narratives, with eight narratives in each noise level). Only neural data from the same sixteen narratives as used in this study were selected for the comparison between the young listeners versus the old listeners.

### Data analysis

#### Preprocessing

A two-step preprocessing method based on HoMER2 software package (*74*) was employed to remove possible motion artifacts. The first step utilized an algorithm grounded in target principal component analysis (function: hmrMotionCorrectPCArecurse; input parameters: tMotion = 0.5, tMask = 1, STDthresh = 30, AMPthresh = 0.5, nSV = 0.97, maxIter = 5). This algorithm discerned artifact-associated principal components and then reconstructed the cleaned signals using the remaining components (*75*). Subsequently, the second step adopted a linear interpolation method (*76*), which identified the motion artifacts based on a predetermined threshold (function: hmrMotionArtifactByChannel; input parameters: tMotion = 0.5, tMask = 1, STDEVthresh = 30, AMPthresh = 0.5), and then corrected these artifacts through cubic spline interpolation (function hmrMotionCorrectSpline; input parameters: p = 0.99). The above algorithms and parameters were all in consistence with that for young listeners (*35*).

#### Inter-brain neural coupling analysis

The Wavelet Transform Coherence (WTC) was employed to analyze the relationship between the speaker’s and listener’s neural activity. WTC allows for the assessment of cross-correlation between two physiological signal series across frequency and time dimensions (*77*). Each trial during the task sessions was initially segmented and further extended to a duration of 300 seconds, which included the 90-second trial and two additional 105-second periods both preceding and following the trial. This extension aimed to facilitate a reliable computation of inter-brain couplings across the desired frequency range. WTC was computed over the 300-second fNIRS signal segments from both the listener and the corresponding speaker, resulting in a coherence matrix organized across time and frequency domains. In the time domain, coherence values were averaged across the entire 90-second trial and subsequently converted to Fisher-*z* values. In the frequency domain, 75 frequency bins ranging from 0.01 to 0.7 Hz were selected for subsequent analysis. Frequencies outside this range were excluded to mitigate aliasing of higher-frequency physiological noise, like cardiac activity (∼0.8–2.5 Hz), and very-low-frequency fluctuations (*78*). Next, coherence values from four trials under the same noise levels were averaged. The coherence value for the middle 90 s of the resting-state session were calculated in the same way. The above calculation was conducted for all speaker-listener channel combinations, creating a total of 1296 cross-channel combinations (36 channels from the speaker × 36 channels from the listener). Consequently, each listener had coherence value under five different conditions, i.e., four speech conditions vs. the resting-state condition, across 1296 channel combinations and 75 frequency bins (0.01-0.07 Hz).

At the group level, a repeated-measures analysis of variance (rmANOVA) was applied to examine the modulation of speech condition on the speaker-listener neural coupling for each channel combination and frequency bin. A nonparametric cluster-based permutation method was utilized to address the issue of multiple comparisons (*79*). This method grouped neighboring frequency bins with uncorrected p-values below 0.05 into clusters and summed the *F*-statistics of these bins as the cluster statistic. A null distribution was generated with the maximum cluster-level statistics from 1,000 permutations. In each permutation, the clusters were calculated with labels of different trials in five conditions shuffled within each participant. Corrected *p*-values for each cluster were derived by comparing the cluster-level statistics to the null distribution. A cluster was significant if the *p*-value was lower than 0.05 level. For each cluster, the speaker-listener neural coupling within the corresponding frequency bins were averaged and subjected to further analysis.

#### Comparison between old listeners and young listeners

A two-factor mixed-design ANOVA (within-subject factor: noise level; between-subject factor: listener group) was used to compare the speaker-listener neural coupling between the old listeners and young listeners. Post hoc analysis was further performed whether the main effect of listener group or the interaction effect between noise level and listener group was significant.

Except for the clusters obtained from the old listeners, the clusters showing significant speaker-listener neural coupling in the young listeners was also included into analysis. These clusters centered over the listener’s left IFG (CH17), right MTG and IPL (CH35-36). They covered distributed brain areas at the speaker’s side, including the bilateral prefrontal cortex (CH4-7/9/11-12/14-15), bilateral postCG (CH22/32) and bilateral IPL (CH23-24/26/30/32-34). The brain regions of these channel combinations are shown in Figure 2A. The channel combination and the frequency band of these clusters are listed in Table S2.

#### Analysis of behavioral relevance

To assess the behavioral relevance of the speaker-listener neural coupling, the Spearman correlation was calculated between the coherence values and old listeners’ speech comprehension performance at each noise level.

#### Single-brain activation analysis

To better explain the mechanism of those brain regions showing significant speaker-listener neural coupling, the neural activity of the brain regions was further analyzed. The preprocessed fNIRS signals were first filtered to 0.01-0.03Hz, which was determined by the above speaker-listener neural coupling analysis. Next, they were normalized by the mean value and standard deviation from the middle 90 s of the resting-state session, and then averaged over the entire 90-s duration for each trial. Then, they were averaged among trials under the same noise level, and further averaged among multiple channels within the same region or network, i.e., prefrontal cortex, ventral language regions or dorsal language regions. Lastly, at the group level, the significance and the behavioral relevance of these neural activation were respectively examined by *t*-test and Spearman correlation analysis.

## Author contributions

Conceptualization: LZR, LY, ZXM, ZD, WS; Methodology: LZR, LY, ZXM, KNN, ZXY, JZR, ZD, WS; Investigation: LZR, LY, ZXM, KNN, ZXY, JZR; Visualization: LZR, ZD; Supervision: AKE, ZD, WS; Writing—original draft: LZR, LY, ZD, WS; Writing—review & editing: LZR, LY, ZXM, AKE, ZD, WS.

## Data and materials availability

All data needed to evaluate the conclusions in the paper are present in the paper and/or the Supplementary Materials. Data supporting this study are available at https://doi.org/10.5061/dryad.sf7m0cgdn

