## Supplemental Materials for "Compensatory Mechanisms for Preserving Speech-in-Noise Comprehension Involve Prefrontal Cortex in Older Adults"

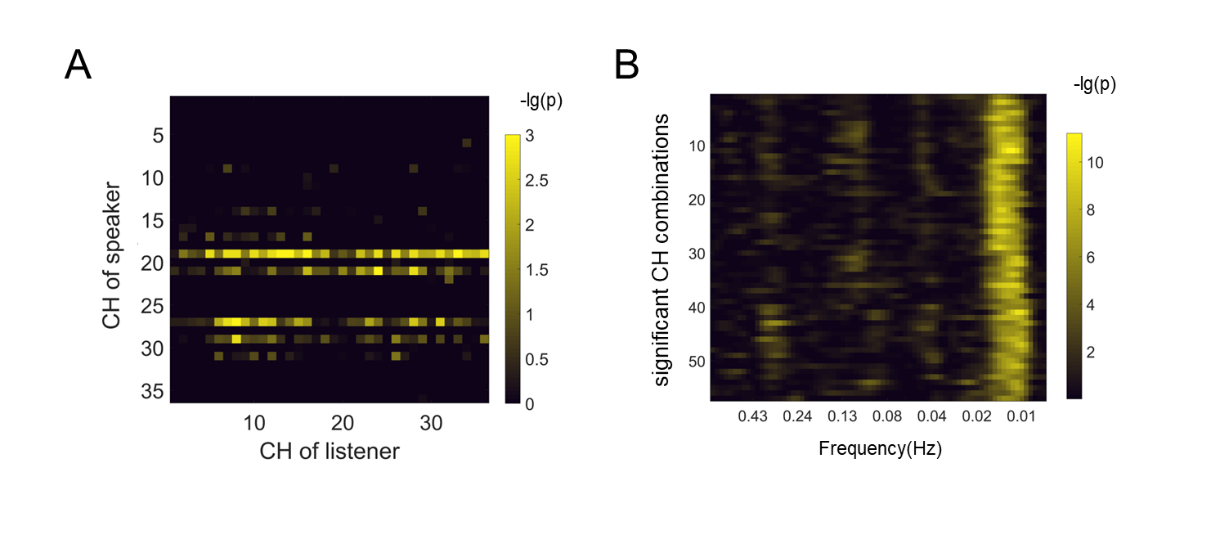

Figure S1. Results of speaker-listener neural coupling analysis. (A) the significance the maximum cluster of each channel combinations. (B) The significance for all frequency bins from 0.01-0.7 Hz of the significant clusters. The frequency band for those clusters is 0.01-0.03Hz.

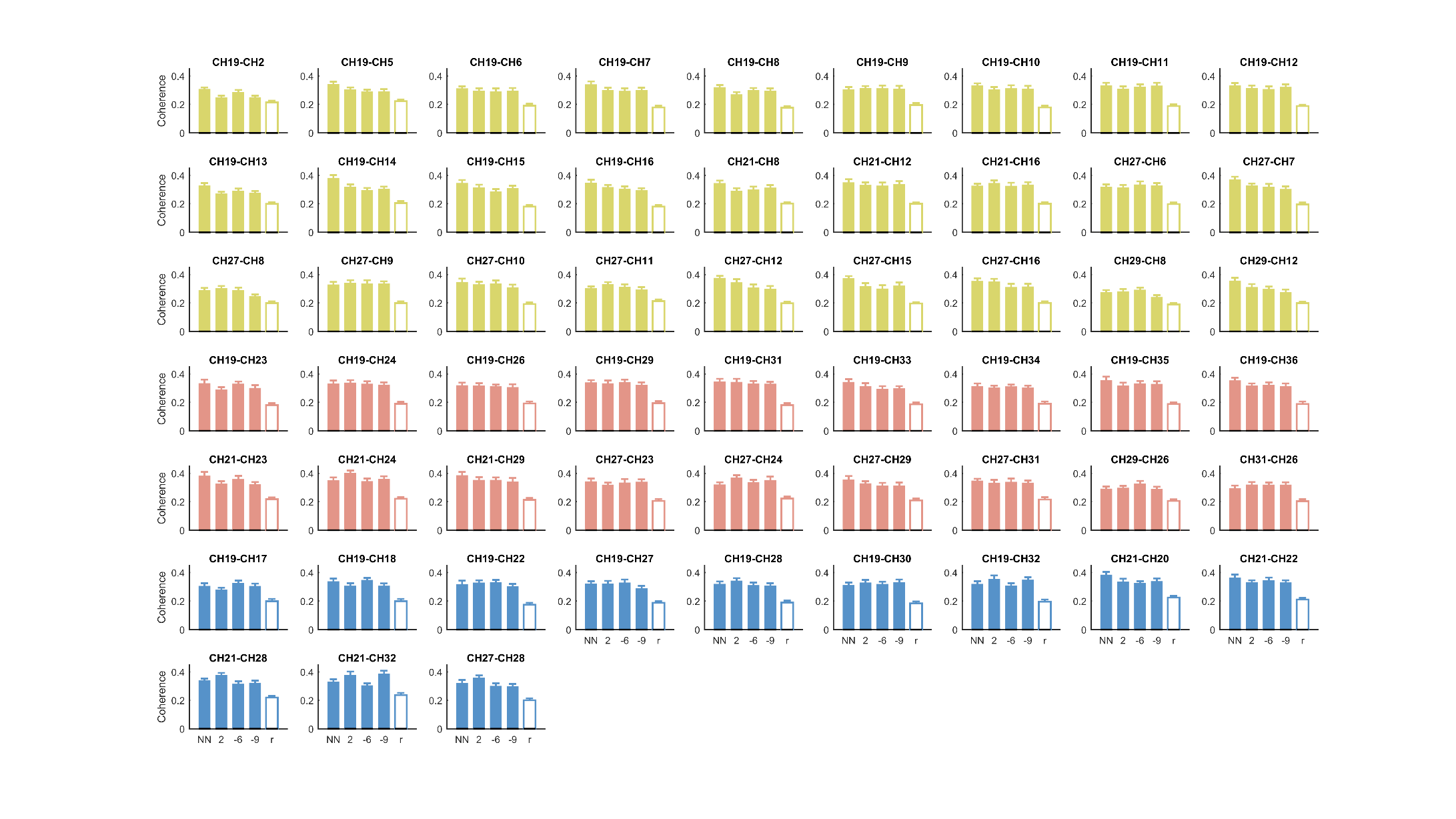

Figure S2. Speaker-listener neural coupling for all clusters. Error bar means the standard error for each condition. Three colors represent the clusters from the old listener’s prefrontal cortex (yellow), ventral brain regions (red), and dorsal language regions (blue).

NN = no noise; r = resting-state condition.

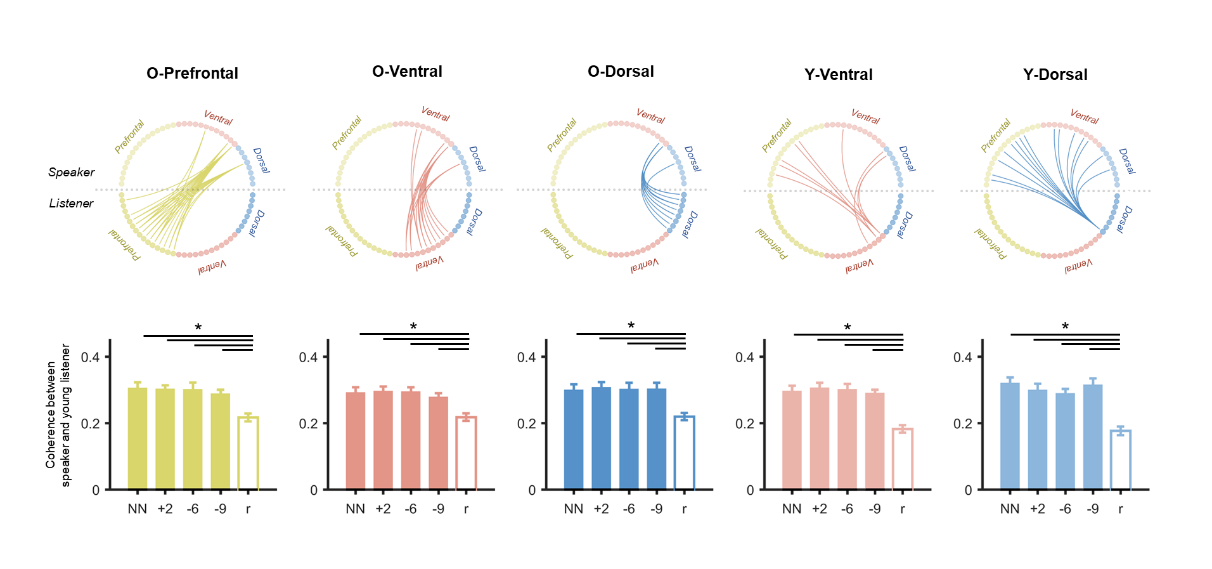

Figure S3. The young listener’s speaker-listener neural coupling at various inter-brain neural patterns. For all five patterns, the speaker-listener neural coupling was higher in four noise levels than the resting-state baseline (*p*s < .05, FDR corrected). Error bar means the standard error for each condition.

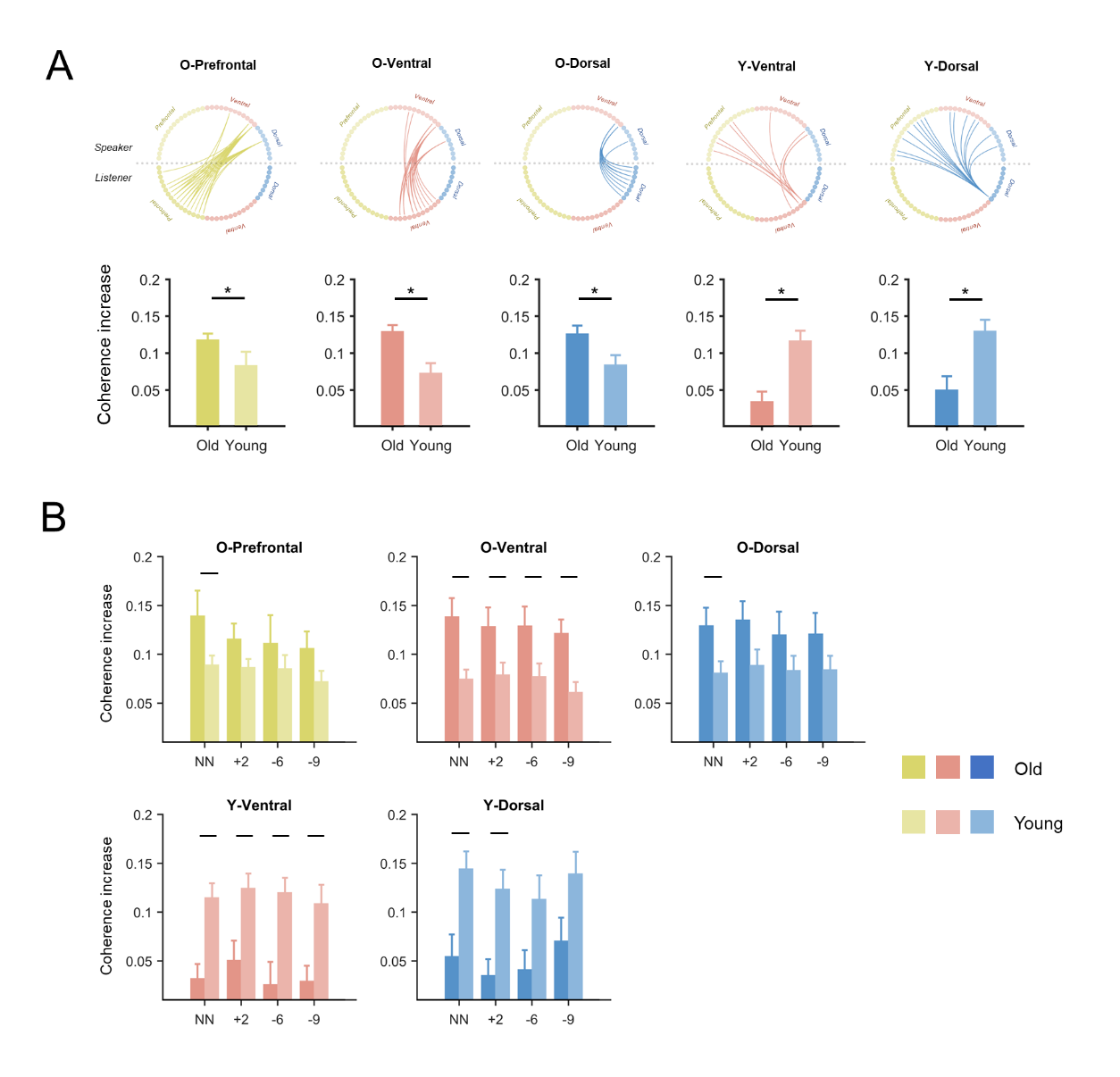

Figure S4. Comparison of speaker-listener neural coupling between old listeners and young listeners. (A) Comparison of neural coupling averaged across noise levels. The coherence from the baseline was first subtracted from the four noise levels and then averaged. The comparison in all five inter-brain neural patterns were significant (*p*s < .05, FDR corrected). For the O-Prefrontal, O-Ventral and O-Dorsal patterns, the coherence of the old listeners was higher than young listeners. In contrast, for the Y-Ventral and Y-Dorsal patterns, the coherence of young listeners was higher than old listeners. (B) Comparison of neural coupling under each noise level. Significant results of comparison were marked by black lines (*p*s < .05, FDR corrected). Error bar means the standard error for each condition.

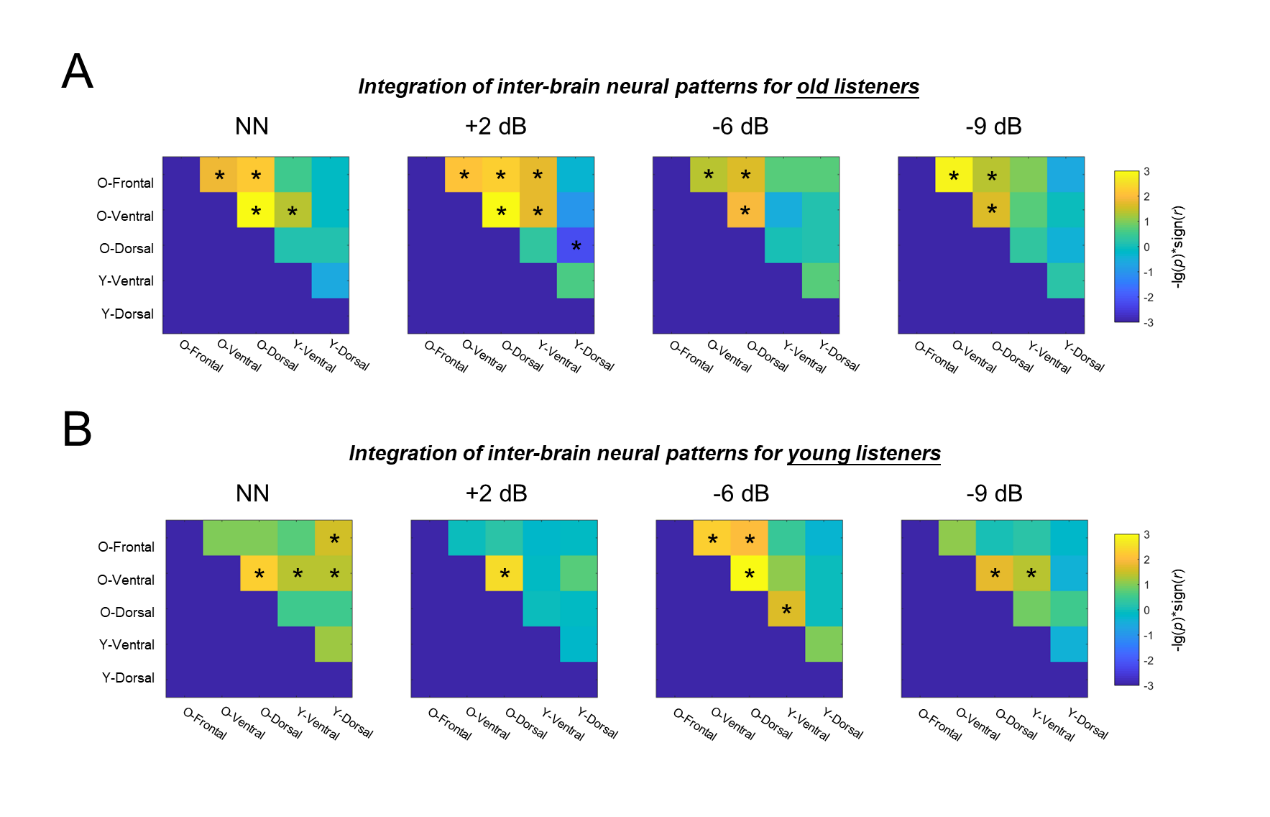

Fig. S5. Correlation matrix for all inter-brain neural patterns for the old listeners (A) and the young listeners (B). Significant correlation coefficients were marked by the asterisks (*p* < .05, FDR corrected).

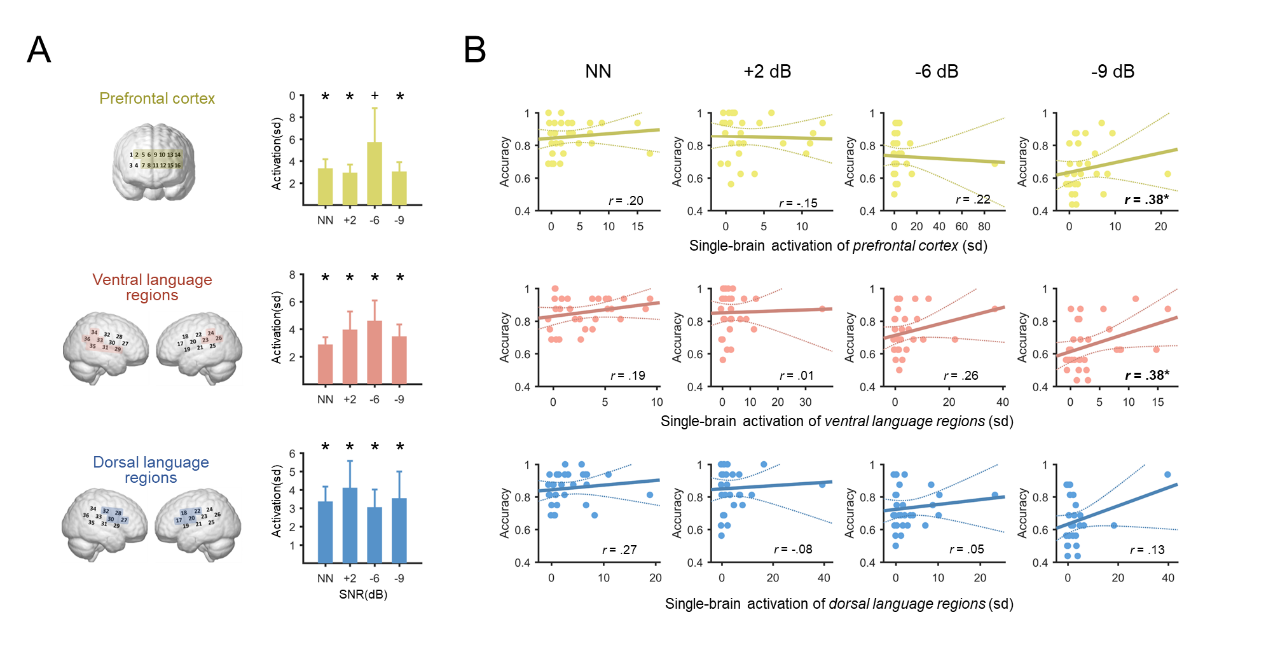

Fig. S6. Single-brain neural activation of the old listeners. (A) Neural activation of these brain regions showing significant speaker-listener neural coupling. The activation of the prefrontal cortex was significant at NN, +2 dB and -9 dB, and marginally significant at -6 dB. The ventral and dorsal language regions were all significantly activated in all four conditions. The error bar means the standard error. The asterisk (*) means significant result (*p* < .05 after FDR correction); the cross (+) means marginal significant result (*p* < .10 after FDR correction). (B) Correlation between the comprehension score and activation of each inter-brain neural pattern. The activation of the prefrontal cortex and the ventral language regions were positively correlated with comprehension score at -9 dB (Spearman *r*s = .38, .38; *p*s = .041, .045, uncorrected). The other correlation coefficients were not significant.

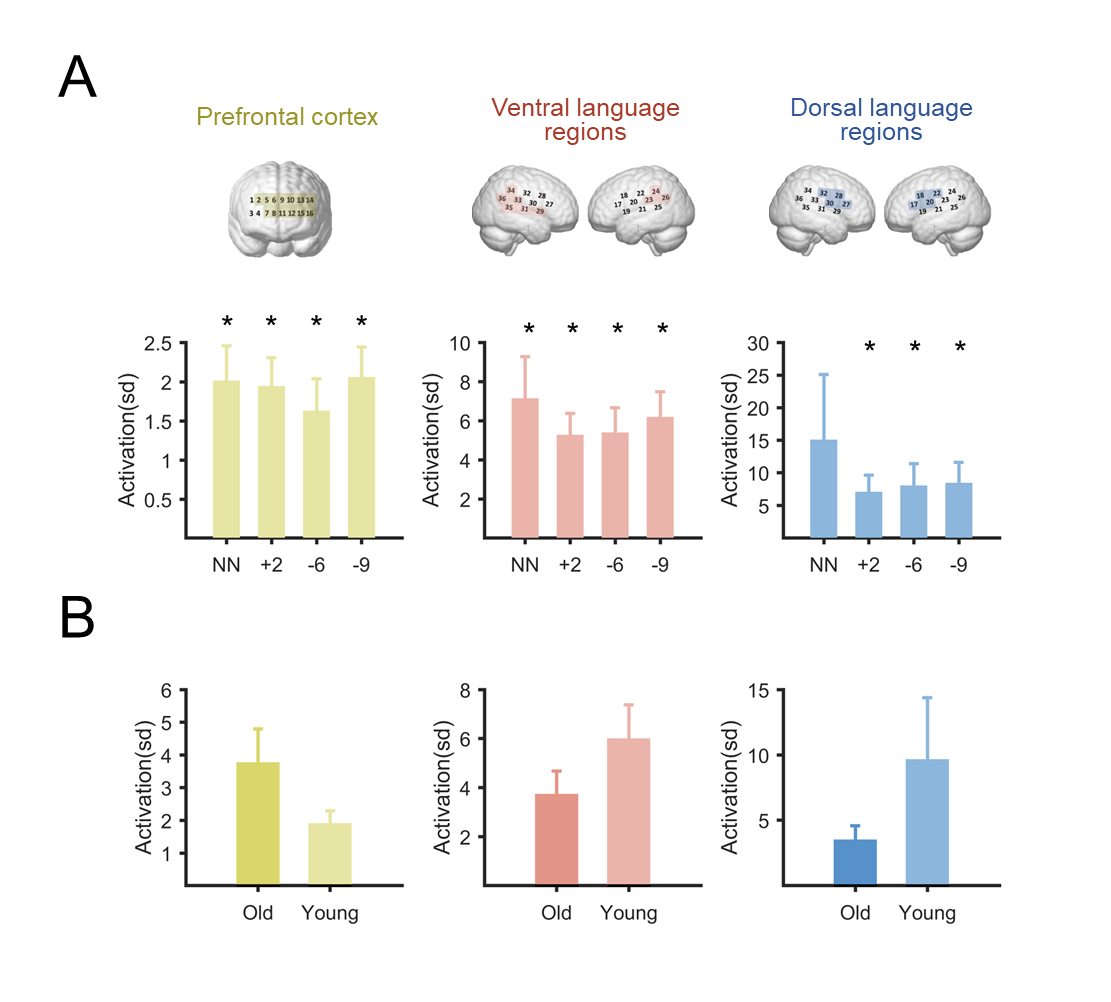

Figure S7. (A) Single-brain neural activation of young listeners. The neural activation of prefrontal cortex, ventral language regions and dorsal language regions were significant in all four noise levels (*p*s < .05, FDR corrected), except for the dorsal language regions in NN (*p* = .15). (B) Comparison of neural activation between old listeners and young listeners. For the prefrontal cortex, the group average of old listeners was higher than the young listeners; while for the ventral and dorsal language regions, the average of young listeners was higher. However, these comparisons were not statistically significant (*p*s = .21, .17, .10).

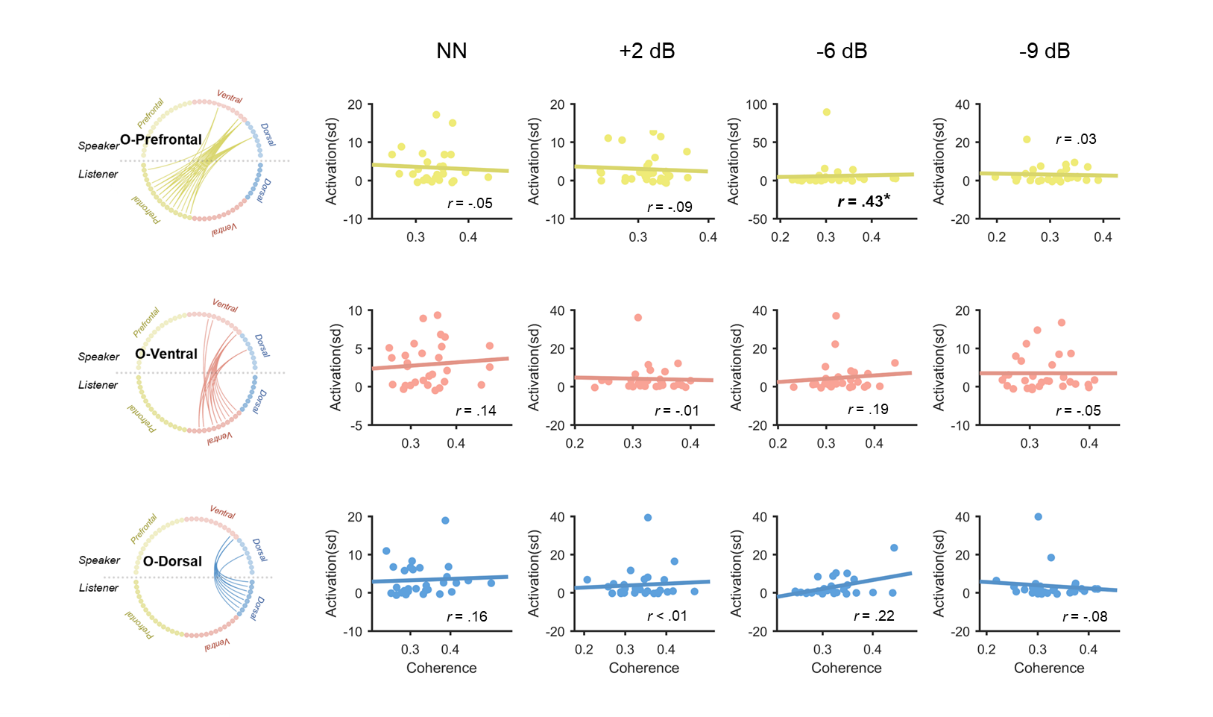

Figure S8. Correlation between the old listener’s single-brain activation and their neural coupling to the speaker. The only significant correlation was observed in the prefrontal cortex and in -6 dB (Spearman *r* = .43, *p* = .019, uncorrected). This correlation remained significant after removing the large outlier point (Spearman *r* = .44, *p* = .019, uncorrected). The other correlation coefficients were non-significant (*p*s > .17).

Table S1. Significant clusters in old adults.

| **CH of speaker** | **CH of listener** | **Frequency range** | **Cluster Significance/p** |
| --- | --- | --- | --- |
| 19 | 2 | 0.01-0.03Hz | 0.038 |
| 19 | 5 | 0.01-0.03Hz | 0.002 |
| 19 | 6 | 0.01-0.02Hz | 0.045 |
| 19 | 7 | 0.01-0.02Hz | 0.002 |
| 19 | 8 | 0.01-0.03Hz | 0.002 |
| 19 | 9 | 0.01-0.02Hz | 0.034 |
| 19 | 10 | 0.01-0.02Hz | 0.002 |
| 19 | 11 | 0.01-0.02Hz | 0.009 |
| 19 | 12 | 0.01-0.02Hz | 0.002 |
| 19 | 13 | 0.01-0.03Hz | 0.001 |
| 19 | 14 | 0.01-0.02Hz | 0.001 |
| 19 | 15 | 0.01-0.02Hz | 0.005 |
| 19 | 16 | 0.01-0.02Hz | 0.001 |
| 19 | 17 | 0.01-0.02Hz | 0.026 |
| 19 | 18 | 0.01-0.02Hz | 0.006 |
| 19 | 22 | 0.01-0.02Hz | 0.005 |
| 19 | 23 | 0.01-0.02Hz | 0.032 |
| 19 | 24 | 0.01-0.02Hz | 0.002 |
| 19 | 26 | 0.01-0.02Hz | 0.002 |
| 19 | 27 | 0.01-0.02Hz | 0.026 |
| 19 | 28 | 0.01-0.02Hz | 0.002 |
| 19 | 29 | 0.01-0.02Hz | 0.009 |
| 19 | 30 | 0.01-0.02Hz | 0.008 |
| 19 | 31 | 0.01-0.02Hz | 0.002 |
| 19 | 32 | 0.01-0.02Hz | 0.010 |
| 19 | 33 | 0.01-0.03Hz | 0.001 |
| 19 | 34 | 0.01-0.02Hz | 0.005 |
| 19 | 35 | 0.01-0.02Hz | 0.006 |
| 19 | 36 | 0.01-0.02Hz | 0.002 |
| 21 | 8 | 0.01-0.02Hz | 0.027 |
| 21 | 12 | 0.01-0.02Hz | 0.041 |
| 21 | 16 | 0.01-0.02Hz | 0.006 |
| 21 | 20 | 0.01-0.02Hz | 0.023 |
| 21 | 22 | 0.01-0.02Hz | 0.010 |
| 21 | 23 | 0.01-0.02Hz | 0.038 |
| 21 | 24 | 0.01-0.02Hz | 0.001 |
| 21 | 28 | 0.01-0.02Hz | 0.002 |
| 21 | 29 | 0.01-0.02Hz | 0.029 |
| 21 | 32 | 0.01-0.02Hz | 0.041 |
| 27 | 6 | 0.01-0.02Hz | 0.008 |
| 27 | 7 | 0.01-0.02Hz | 0.002 |
| 27 | 8 | 0.01-0.03Hz | <0.001 |
| 27 | 9 | 0.01-0.02Hz | 0.006 |
| 27 | 10 | 0.01-0.02Hz | 0.045 |
| 27 | 11 | 0.01-0.03Hz | 0.003 |
| 27 | 12 | 0.01-0.02Hz | 0.005 |
| 27 | 15 | 0.01-0.02Hz | 0.009 |
| 27 | 16 | 0.01-0.02Hz | 0.019 |
| 27 | 23 | 0.01-0.02Hz | 0.010 |
| 27 | 24 | 0.01-0.02Hz | 0.045 |
| 27 | 28 | 0.01-0.02Hz | 0.004 |
| 27 | 29 | 0.01-0.02Hz | 0.039 |
| 27 | 31 | 0.01-0.02Hz | 0.003 |
| 29 | 8 | 0.01-0.03Hz | 0.002 |
| 29 | 12 | 0.01-0.02Hz | 0.046 |
| 29 | 26 | 0.01-0.03Hz | 0.032 |
| 31 | 26 | 0.01-0.03Hz | 0.041 |

Table S2. Significant clusters in young adults.

| **CH of speaker** | **CH of listener** | **Frequency range** |
| --- | --- | --- |
| 4 | 17 | 0.01-0.03Hz |
| 5 | 17 | 0.01-0.03Hz |
| 6 | 17 | 0.01-0.03Hz |
| 7 | 17 | 0.01-0.03Hz |
| 9 | 17 | 0.01-0.03Hz |
| 11 | 17 | 0.01-0.02Hz |
| 14 | 17 | 0.01-0.03Hz |
| 15 | 17 | 0.01-0.03Hz |
| 22 | 17 | 0.01-0.03Hz |
| 23 | 17 | 0.01-0.03Hz |
| 24 | 17 | 0.01-0.03Hz |
| 26 | 17 | 0.01-0.03Hz |
| 30 | 17 | 0.01-0.03Hz |
| 33 | 17 | 0.01-0.03Hz |
| 34 | 17 | 0.01-0.03Hz |
| 32 | 33 | 0.01-0.02Hz |
| 7 | 35 | 0.01-0.03Hz |
| 14 | 35 | 0.01-0.03Hz |
| 30 | 35 | 0.01-0.03Hz |
| 6 | 36 | 0.01-0.03Hz |
| 11 | 36 | 0.01-0.02Hz |
| 12 | 36 | 0.01-0.03Hz |
| 30 | 36 | 0.01-0.03Hz |
| 33 | 36 | 0.01-0.03Hz |

Table S3. MNI coordinates and anatomical labels for each fNIRS channel.

| **Channel** | **MNI(x,y,z)** | | | **Anatomical label and percentage of overlap** |
| --- | --- | --- | --- | --- |
| CH1 | 52.67 | 56.67 | 34.67 | R middle frontal gyrus, 0.8528  R inferior frontal gyrus, 0.1472 |
| CH2 | 40.00 | 64.67 | 41.33 | R middle frontal gyrus, 1 |
| CH3 | 54.00 | 67.67 | 1.67 | R middle frontal gyrus, 0.1098  R inferior frontal gyrus, 0.8902 |
| CH4 | 41.33 | 75.67 | 6.67 | R middle frontal gyrus, 0.8121  R inferior frontal gyrus, 0.1879 |
| CH5 | 25.67 | 70.67 | 45.00 | R superior frontal gyrus, 0.0539  R middle frontal gyrus, 0.9461 |
| CH6 | 9.00 | 74.00 | 45.00 | L superior frontal gyrus, 0.0004  R superior frontal gyrus, 0.6160  R middle frontal gyrus, 0.3836 |
| CH7 | 26.33 | 83.00 | 11.00 | R superior frontal gyrus, 0.0130  R middle frontal gyrus, 0.9870 |
| CH8 | 8.67 | 87.00 | 10.67 | L superior frontal gyrus, 0.0203  R superior frontal gyrus, 0.4915  R middle frontal gyrus, 0.4881 |
| CH9 | -7.33 | 72.67 | 47.67 | L superior frontal gyrus, 0.7198  R superior frontal gyrus, 0.1829  L middle frontal gyrus, 0.0973 |
| CH10 | -22.33 | 69.67 | 46.00 | L superior frontal gyrus, 0.3527  L middle frontal gyrus, 0.6473 |
| CH11 | -9.67 | 86.00 | 15.00 | L superior frontal gyrus, 0.8082  R superior frontal gyrus, 0.0616  L middle frontal gyrus, 0.1301 |
| CH12 | -26.00 | 82.33 | 13.00 | L superior frontal gyrus, 0.1812  L middle frontal gyrus, 0.8188 |
| CH13 | -36.00 | 63.67 | 42.67 | L middle frontal gyrus, 1 |
| CH14 | -49.00 | 55.00 | 37.67 | L middle frontal gyrus, 1 |
| CH15 | -39.00 | 75.33 | 9.33 | L middle frontal gyrus, 0.9211  L inferior frontal gyrus, 0.0789 |
| CH16 | -52.00 | 66.67 | 5.33 | L middle frontal gyrus, 0.4538  L inferior frontal gyrus, 0.5462 |
| CH17 | -74.00 | 26.33 | 4.33 | L inferior frontal gyrus, 0.9799  L precentral gyrus, 0.0201 |
| CH18 | -74.00 | 14.67 | 30.33 | L middle frontal gyrus, 0.0469  L inferior frontal gyrus, 0.3321  L precentral gyrus, 0.6209 |
| CH19 | -79.00 | 7.67 | -9.67 | L superior temporal gyrus, 0.5567  L middle temporal gyrus, 0.4433 |
| CH20 | -81.00 | -3.67 | 16.33 | L precentral gyrus, 0.1632  L postcentral gyrus, 0.8090  L supramarginal gyrus, 0.2243  L superior temporal gyrus, 0.0035 |
| CH21 | -85.00 | -22.33 | 1.33 | L superior temporal gyrus, 0.4528  L middle temporal gyrus, 0.5472 |
| CH22 | -77.67 | -15.00 | 42.67 | L postcentral gyrus, 0.4642  L supramarginal gyrus, 0.5358 |
| CH23 | -82.00 | -36.00 | 28.00 | L supramarginal gyrus, 0.7065  L superior temporal gyrus, 0.2935 |
| CH24 | -75.00 | -47.67 | 51.00 | L supramarginal gyrus, 0.5133  L angular gyrus, 0.4867 |
| CH25 | -80.00 | -55.00 | 9.00 | L supramarginal gyrus, 0.0033  L angular gyrus, 0.0327  L superior temporal gyrus, 0.3300  L middle temporal gyrus, 0.6340 |
| CH26 | -74.00 | -65.67 | 32.67 | L angular gyrus, 0.9963  L middle occipital gyrus, 0.0037 |
| CH27 | 75.00 | 28.00 | 0.00 | R inferior frontal gyrus, 0.8436  R precentral gyrus, 0.0977  R superior temporal gyrus, 0.0586 |
| CH28 | 76.00 | 18.67 | 26.67 | R inferior frontal gyrus, 0.1070  R precentral gyrus, 0.8930 |
| CH29 | 81.00 | 7.67 | -14.33 | R superior temporal gyrus, 0.3322  R middle temporal gyrus, 0.6678 |
| CH30 | 83.00 | -1.33 | 11.67 | R precentral gyrus, 0.0190  R postcentral gyrus, 0.7911  R superior temporal gyrus, 0.1899 |
| CH31 | 87.00 | -22.33 | -3.67 | R superior temporal gyrus, 0.2866  R middle temporal gyrus, 0.7134 |
| CH32 | 81.00 | -13.00 | 37.67 | R postcentral gyrus, 0.4071  R supramarginal gyrus, 0.5929 |
| CH33 | 85.00 | -33.67 | 20.67 | R supramarginal gyrus, 0.3823  R angular gyrus, 0.1223  R superior temporal gyrus, 0.4954 |
| CH34 | 79.00 | -46.00 | 44.00 | R supramarginal gyrus, 0.5000  R angular gyrus, 0.5000 |
| CH35 | 81.00 | -56.00 | 3.00 | R middle temporal gyrus, 1 |
| CH36 | 75.33 | -66.33 | 25.33 | R angular gyrus, 0.8877  R middle occipital gyrus, 0.0737  R middle temporal gyrus, 0.0386 |
